# Reference Point-Dependent Reinforcement Learning in Humans and Rats

**DOI:** 10.1101/2025.04.11.648377

**Authors:** Lachlan Ferguson, Magdalena Soukupova, Sebastien Bouret, Stefano Palminteri, Shauna Parkes

**Affiliations:** Univ. Bordeaux, CNRS, INCIA, UMR 5287, F-33000 Bordeaux, France; Laboratoire des Neurosciences Cognitives et Computationnelles, Institut National de la Santé et de la Recherche Médicale, Paris, France; Département d’Etudes Cognitives, Ecole Normale Supérieure, PSL Research University, Paris, France; Motivation Brain and Behaviour team, Brain and Spine Institute, Paris, France

## Abstract

Previous studies indicate that rewards and punishments in reinforcement learning are encoded in a relative manner. Reference point-dependence, a valuation bias shared by eminent adaptation level and prospect theories, is often cited as the underlying computational mechanism. However, whether these behavioural and computational mechanisms are preserved across species is unknown. To fill this gap, we designed a reinforcement learning task that was adapted for both humans *(homo sapiens sapiens)* and rats (*rattus norvegicus*). Behavioural analyses indicated robust relative value encoding in both species. At the computational level, reference point-dependence provided a reliable account of reinforcement learning behaviour in humans and rats. Despite these major similarities between species, some distinct differences in behavioural and computational modelling parameters were also observed. Overall, our study demonstrates that relative value encoding is a robust feature of reinforcement learning that is conserved across human and rat species.

## Introduction

Economic decision-making is not always constrained by rational choice axioms (Simon, 1955; von Neumann & Morgenstern, 1944). Rather, choices appear to be context-dependent, in that their subjective values are influenced by the presence of other options presented simultaneously or in the recent past (Helson, 1948; Kahneman, 2003; Kahneman & Tversky, 1979; Parducci, 1965; Simon, 1955). In human decision making, such contextual influences have been demonstrated in a large variety of tasks, ranging from psychophysical perceptual judgements, decisions among lotteries and, more recently, outcome representation in reinforcement learning (Helson, 1948; Louie & Glimcher, 2012; Palminteri & Lebreton, 2021; Parducci, 1965; Rangel & Clithero, 2012).

In reinforcement learning (RL), a well-documented consequence of context-dependent outcome valuation is the occurrence of suboptimal (i.e., reward minimizing) choices, whereby established relative values in prior contexts conflict with optimal (i.e., reward maximising) choices in new contexts (Bavard et al., 2018; Hayes & Wedell, 2023; Klein et al., 2017; Palminteri et al., 2015). In humans, such studies typically implement a two-phase protocol that consists of an initial learning phase, where pairs of options—say, AB or CD—are always presented together, followed by a transfer phase, where options are rearranged into novel pairs—say, BC. The pattern of choices during the transfer phase reveals the extent to which context-dependent learning drives behaviour, particularly when optimal choices based on absolute and relative values diverge (Palminteri & Lebreton, 2021).

Context-dependent RL appears to be a wide-spread feature of human cognition that emerges cross-culturally (Anlló et al., 2024), indicating that it is a reliable cognitive feature in our species. Moreover, evidence for context-dependent RL has also been observed in other species (e.g., starlings, pigeons, and bumblebees) using behavioural tasks similar to those used in humans (Belke, 1992; Freidin & Kacelnik, 2011; Gibbon, 1995; Monteiro et al., 2013; Pompilio & Kacelnik, 2010; Solvi et al., 2022; Vasconcelos et al., 2013). However, despite their superficial similarity, there has been no direct concomitant assessment across human and non-human choice behaviour using the same task. Indeed, the species-specific idiosyncrasies of reinforcement schedules used in these studies have made it challenging to conduct post hoc comparisons across species, and the computational form of context-dependency in RL in non-human subjects remains unknown.

We reasoned that, if the behavioural options and reinforcement structure were held constant across species, we could more effectively observe similar behaviour and computations in humans and rats. In order to examine context-dependent learning in both these species, we used a structurally similar behavioural task with identical visual cues and reinforcement structures, and compared the same computational models in both species. This allowed us to minimise interspecies variance and enhance the comparability of RL models between species. Adopting this approach makes it possible to determine if context-dependent RL processes are conserved at the mechanistic level, rather than simply reflecting outwardly similar behavioural signatures.

Our results demonstrate that, when faced with options whose values were acquired in different learning contexts, humans and rats prefer contextually high relative value options even if this entails an economic cost. At the computational level, choice behaviour in both species was better explained by a model assuming that rewards are encoded relative to a dynamically learned reference point (Hayes & Wedell, 2022; Palminteri et al., 2015). Despite these major similarities, we also observed some distinctive species-specific behavioural and computational characteristics.

## Results

### Context-dependent learning and transfer in human and rats

Both species performed a reinforcement learning bandit task with two phases: a learning phase and a transfer phase. This design was inspired by previous studies on context-dependent reinforcement learning (Palminteri & Lebreton, 2021; Vasconcelos et al., 2013) and was optimized to minimize differences across species (see **Methods** for further details). During the learning phase, humans and rats were presented with 4 differentially rewarded options (A, B, C, or D) displayed on a screen in stable binary choice contexts (AB or CD contexts) (**Figure 1A-B—Learning phase**). Each option was associated with a specific fixed probability of earning reward (monetary rewards for humans and food rewards for rats), which was adjusted across experiments to explore the effects of different reward probability ratios on context-dependent choice (**Figure 1C**). During the learning phase, contexts were classified as a function of their schedules of reinforcement: a “rich” context (AB) with an overall high reward rate, and a “poor” context (CD) with an overall lower reward rate. Crucially, rich and poor contexts were matched in terms of their objective difficulty (i.e., equal difference in reward probability between options in each context). Humans were given equal information about option values through full feedback, such that they were presented with the outcome for the chosen and unchosen options. Similarly, to avoid a sampling bias in rats, they were given an equal number of forced choice trials for all options, interleaved with unrewarded choice trials.

**Figure 1:**
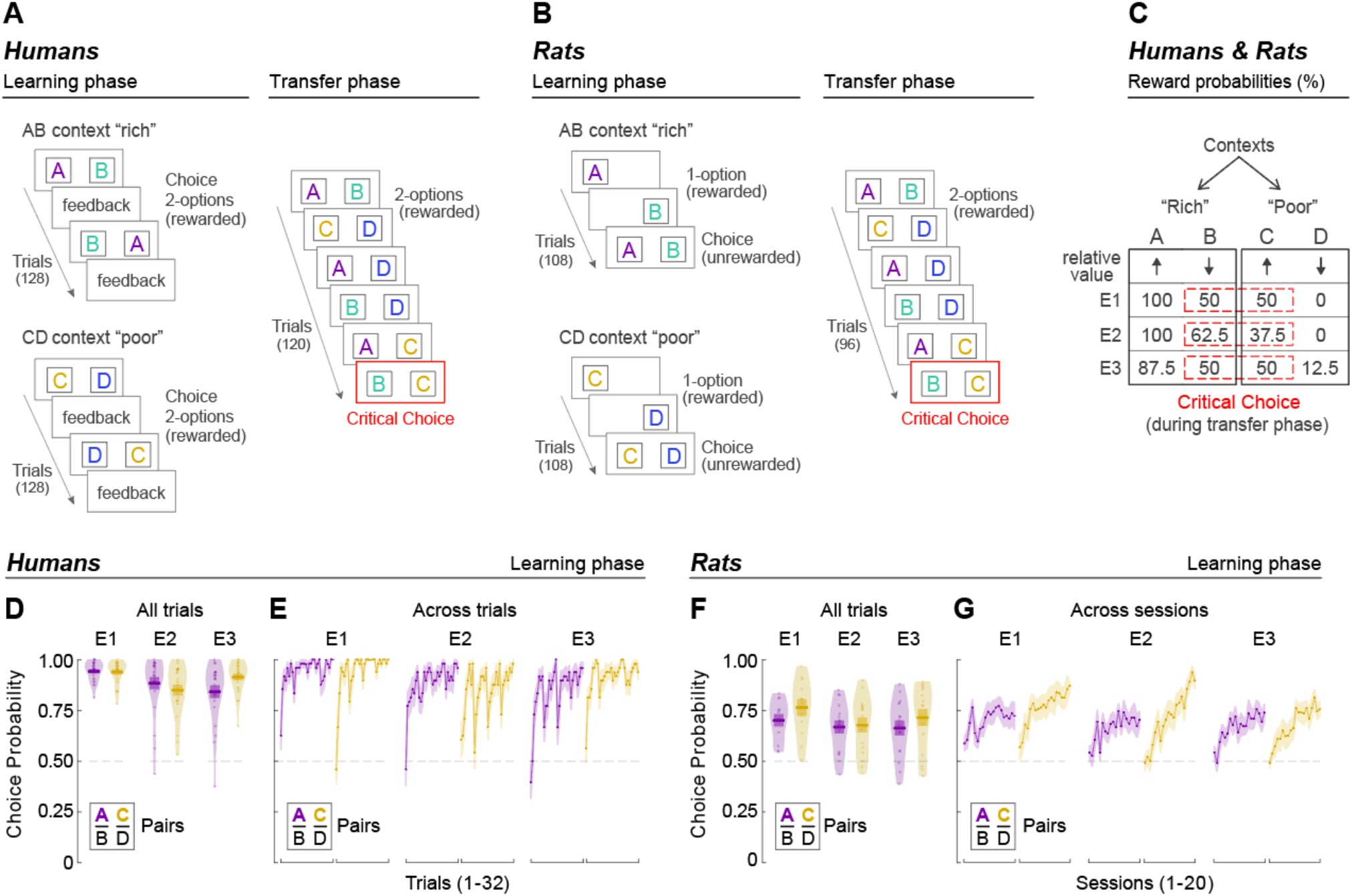
Behavioural protocol and learning phase results. (**A**) and (**B**) task design in humans, and rats respectively. Note that for illustration in the figure options are represented as letters, but they were actually highly discriminable colored gabor patches (see methods). In (A), (B), (C) “critical choice” designates the transfer phase choice which is particularly diagnostic of an absolute or relative learning code. (**C**) Reward probabilities across the three experiments. Note, the number of sessions differed between species. Humans completed the learning and transfer phases over a single session, whereas rats underwent 20 sessions per context during learning, and 10 sessions during the transfer phase (**D**) Accuracy in the learning phase in humans, averaged across all trials. (**E**) Accuracy in the learning phase in humans at the trial-level. (**F**) and (**G**) same as (D) and (E), but for rats. In (D) and (F) individual data points on violin plots represent the average performance per subject. E1, E2 and E3 designate Experiment 1, Experiment 2 and Experiment 3, respectively.

To test for context-dependencies formed in rich and poor environments during learning, options were then presented under rewarded (partial feedback) conditions in all possible binary combinations (AB, CD, AD, BD, AC, BC) during the transfer phase (1 session for humans, 10 sessions for rats) (**Figure 1A-B—Transfer phase**). By examining choices from these pair combinations, we could test if humans and rats encoded information in a context-dependent manner. Specifically, we assigned distinct reward probabilities across three experiments to test BC choices under several unique conditions (**Figure 1C**): in Experiment 1 (E1), options B and C had equivalent reward probabilities but opposing relative values; in Experiment 2 (E2), options B and C had both opposing reward probabilities and relative values; and in Experiment 3 (E3), as in E1, options B and C had equivalent reward probabilities but different relative values—however, the reward probabilities for options A and D were non-deterministic to control for potential reward certainty effects (Tversky & Kahneman, 1986).

### Humans and rats select high value options in rich and poor learning contexts

During the learning phase, the overall probability of choosing the high value options (A in AB, and C in CD) in both “rich” and “poor” contexts was significantly above chance in both species, and this was true for all experiments, regardless of the reward probability ratios (**Figure 1D, 1F**) (all p<0.001, details in **Supplemental Table 1**). A notable difference between the species during the learning phase was the rate at which each species formed their preferences. Humans acquired preferences for high value options rapidly over trials within a single session, whereas rats ‘preference for high value options grew more slowly over several sessions (**Figure 1E, 1G, Supplemental Table 1**). These data indicate that both humans and rats consistently directed their choices towards options associated with higher reward probability, and that humans achieve this at a rate observable at the trial level (within a single session) and rats achieve this at a rate more easily observed at the session level.

### Humans and rats share similar context-dependent choice biases

In Experiment 1 (E1), options B and C had equivalent reward probabilities (50:50), but different relative values (B = low; C = high) in their respective “rich” and “poor” learning contexts. During the transfer phase, when presented with all possible pairs of options, humans and rats chose the high reward probability option in all pairs except the critical BC pair (**Figure 2A-*E1*, 2B-*E1***) (humans: all ≤ p ≤0.010, **Supplemental Table 2**; rats: all p≤0.018, **Supplemental Table 3**). In fact, for both species, during the initial transfer trials, option B was chosen significantly less than chance, indicating a clear preference for the high relative value option C (**Figure 2C-*E1*, 2D-*E1*, Supplemental Figure 1-*E1***; humans: M=0.282, 95%CI [0.217, 0.358], p<0.001 **Supplemental Table 2**; rats: 0.302, 95%CI [0.245, 0.364], p<0.001, **Supplemental Table 3**). The strength of this bias diminished across trials in humans (slope = 0.076, 95% CI [0.017, 0.136], p=0.012, **Supplemental Table 2**), whereas rats ‘preference for the relative high value option remained stable across sessions (**Figure 2C-*E1*, 2D-*E1***, slope = -0.002 95%CI [-0.039, 0.037], p=0.903, **Supplemental Table 3**). Thus, in the transfer phase, both humans and rats bias their preference toward a relative high value option from a “poor” context over a relative low value option of equivalent reward probability from a “rich” context — a bias from which only humans partially recover.

**Figure 2:**
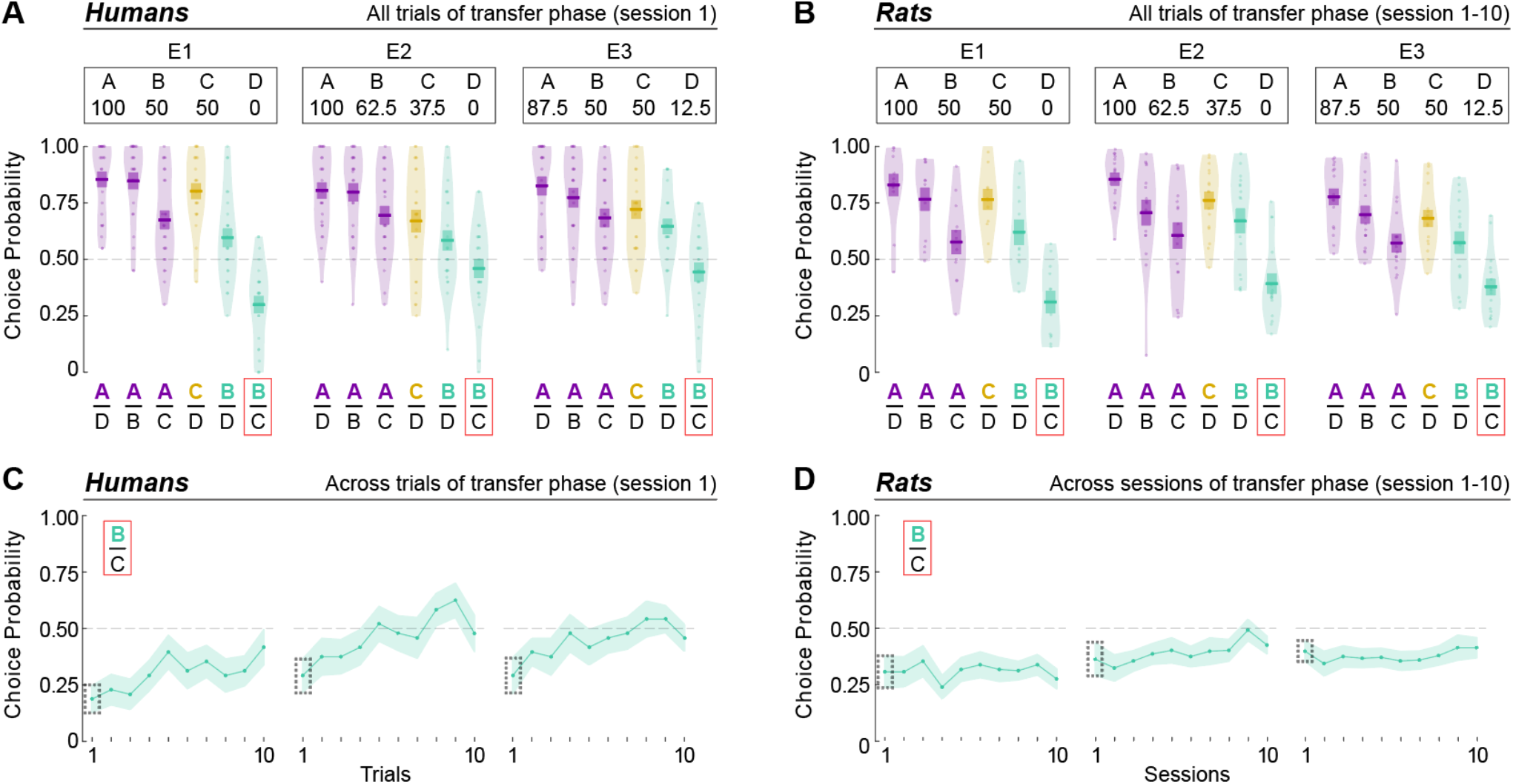
Transfer phase results. Top panels show choice probability for all possible option combinations in the transfer phase, aggregated across all trials (and sessions) in humans (**A**) and rats (**B**) for all three experiments (E1, E2, and E3). Bottom panels present choice probability in the key B|C comparison across trials (humans, **C**) and sessions (rats, **D**). Individual data points represent the average performance per subject (10 trials/pair for humans, and 16 trials/pair/session for rats). In (C) and (D) the dotted black rectangles are used to highlight the first trial (humans) and session (rats) of the BC choice.

Then, we wanted to assess whether this bias towards the option with higher relative value could lead to suboptimal (i.e., reward minimising) choice behaviour. To address this aim, in E2, options B and C had both opposing reward probabilities (B = 62.5%; C = 37.5%) and relative values (B = low; C = high). When presented with all possible pair combinations during the transfer phase, humans and rats once again chose the high reward probability option for all pairs except in the BC pair (**Figure 2A-*E2*, 2B-*E2***, humans: all p≤0.015, rats: all p<0.001, **Supplemental Table 2 and 3**). The overall relative value bias, based on the probability of B choices in BC pairs, was stronger in rats (M=0.387, 95%CI [0.331, 0.446], p<0.001, **Supplemental Table 3**) than in humans (M=0.463, 95%CI [0.38, 0.548], p=0.398, **Table 2**). However, closer inspection shows that humans once again initially display a strong relative value bias (**Figure 2C-*E2*, 2D-*E2*, Supplemental Figure 1A-*E2***) (M=0.272, 95%CI [0.184, 0.382], p<0.001), but partially recover reaching chance-level indifference rapidly across trials (slope =0.101, 95%CI [0.046, 0.155], p<0.001), whereas rats approach chance-level responding only after many sessions (**Figure 2C-*E2*, 2D-*E2***) (slope = 0.053, 95% CI [0.022, 0.083], p<0.001). Thus, the relative values encoded during learning in the “rich” and “poor” contexts initially biased choice, and this suboptimal choice bias found in both species can be partially recovered through re-learning in the transfer context.

Finally, we tested if the context-dependent preferences observed in prior experiments were due to overweighting the value of C when it is paired with a deterministically unrewarded option (option D). In E3, options A or D were now non-deterministic (A = 87.5%; D = 12.5%) and options B and C had equivalent reward probabilities (50:50) but different relative values (B = low; C = high). We found that, much like E1, humans and rats showed a preference for the high reward probability option in all pairs except in the critical BC pair, where option C was significantly preferred (**Figure 2A-*E3*, 2B-*E3***) (humans: all p<0.001, rats: all p≤0.015). Moreover, during the initial transfer trials, humans and rats showed a clear preference for C over B (**Figure 2C-*E3*, 2D-*E3*, Supplemental Figure 1–*E3;*** humans: M=0.299, 95%CI [0.206, 0.411], p<0.001, **Supplemental Table 2**; rats: M=0.358 95%CI [0.301, 0.418], p<0.001, **Supplemental Table 3**). Similarly to E1, the strength of this bias diminished across trials in human subjects (slope=0.073, 95%CI [0.020,0.127], p<0.007, **Supplemental Table 2**), but remained stable across sessions in rats (**Figure 2C-*E3*, 2D-*E3***; slope=0.016, 95%CI [-0.015, 0.046], p=0.312). Thus, the context-dependent preferences that we observed are not simply due to the presence of deterministic options in the learning contexts.

Despite the substantial similarities between humans and rats, a closer inspection of the transfer phase results revealed interesting differences between the two species, particularly regarding the relative values assigned to the extreme options. When examining AC and BD choices (i.e., choices between the two contextually best options and choices between the two contextually worst options), after averaging across experiments and trials, humans demonstrated significantly higher accuracy in the AC compared to the BD contexts (contrast_AB/CD_ = 1.45, z= 4.47, p<0.001). Conversely, rats exhibited significantly greater accuracy when choosing between the two worst options (BD) compared to (AC) (contrast_AB/CD_ = 0.852, z= -4.47, p<0.001). Furthemore, we observed that the difference between AC and BD accuracies were modulated over the course of the transfer phase, generally leading to an increase in accuracy in the AC condition (**Figure 3**). This observation aligns with the accuracy profile observed during the learning phase where we found by numerically higher accuracy in the “poor” compared to the “rich” environment (**Figure 1F** and **1G**).

**Figure 3:**
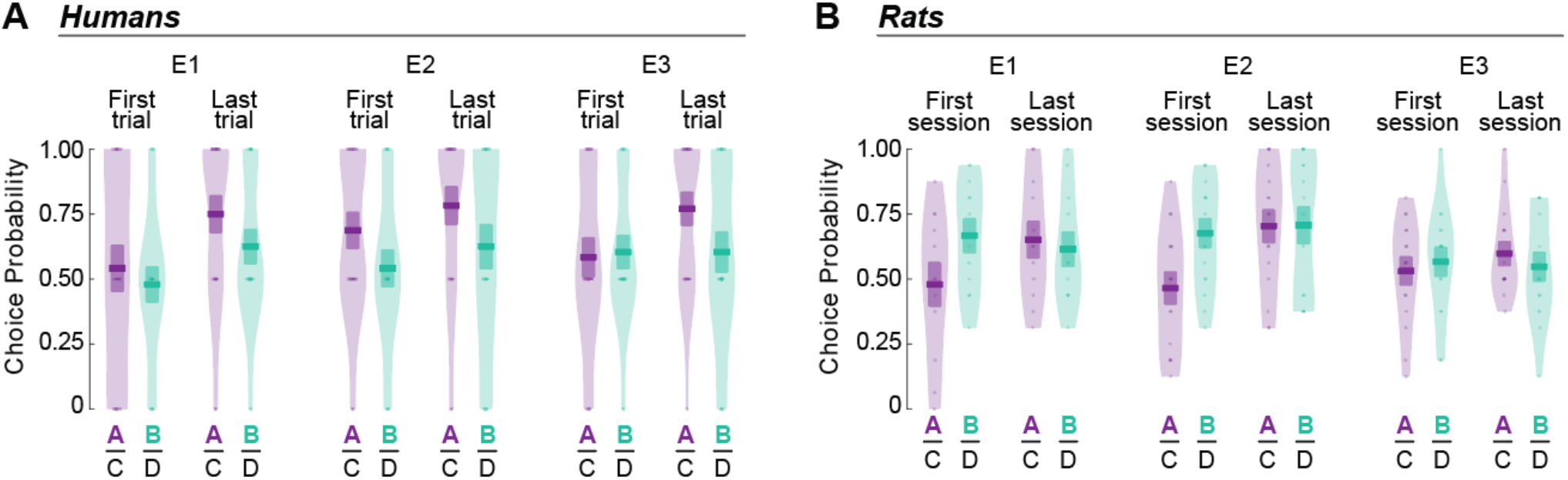
Comparisons of contextually best and worst options at the beginning and end of the transfer phase. Choice probability for the two best options [AC] and the two worst options [BD] in the first and last trials for humans (**A**), and the first and last session for rats (**B**).

### Computational Models

Having found behavioural evidence for relative value encoding in the transfer phase choices (namely preference for C in BC choices), we next turned to computational modelling to assess whether or not the observed behavioural pattern was captured by a known and plausible model of relative value encoding in both species. To do so, we fitted both the learning and transfer phases with different models, which primarily differed in how the choice outcomes were encoded. The “Absolute” model assumes that choice outcomes are encoded in an objective manner. That is, the subjective outcome is equal to the objective outcome.

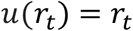

By contrast, in the “Reference” model, the outcome of each option is adjusted as a function of a state-specific, dynamically learned, reference point – which essentially tracks the average value of options ‘ pairs, a process also referred to as “centering” (Naik et al., 2024; Palminteri et al., 2015). Specifically, the “Reference” model assumes that subjects calculate a mean outcome value for each context and subtract it from the actual outcome.

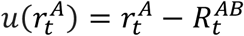

where 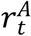 is the objective outcome of option *A* at trial 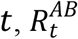 is the reference mean for context *AB* and 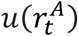 is the normalised outcome (see **Methods** for more details).

Quantitative model comparison indicated that the “Reference” model provided a significantly better fit for both human and rat subjects across all experiments (**Table 1**). To further confirm the validity of the “Reference” model, we simulated data from both the “Absolute” and “Reference” models and compared them to the observed responses in the learning and transfer phases (Palminteri et al., 2017). The “Reference” model, but not the “Absolute” model, was able to replicate the key behavioural patterns, namely, the simulated responses were significantly above chance in the learning phase, and remained above chance for all choice pairs comparisons except the BC pair (**Figure 4**). We also found that, in both species, the learning rate of the learning phase was higher compared to that of the transfer phase (**Table 2**). We also note that the learning rates were much higher in humans compared to rats, even if a formal, statistical, comparison is not possible given differences in the experimental set-up (secondary versus primary reward, complete feedback and number of trials).

**Figure 4.**
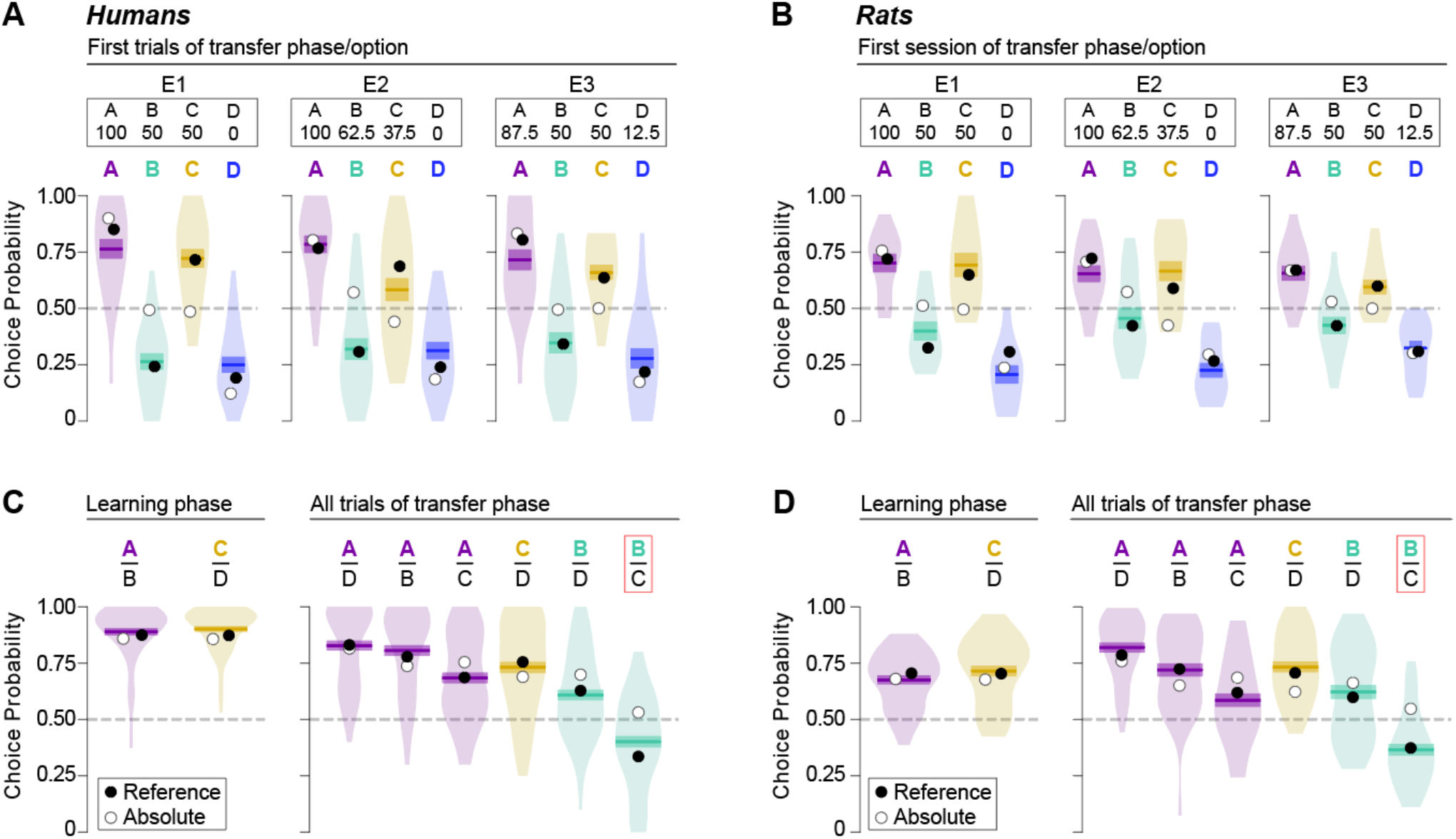
Model falsification for the learning and transfer phases. The reference model (open circles), but not the absolute model (closed circles), was able to replicate the observed patterns of responses (plotted as violin plots) in the transfer test for (**A**) humans in the initial highly context-dependent trials of the transfer phase, irrespective of pairs; and (**B**) for rats in initial highly context-dependent session of the transfer phase, irrespective of pairs. (**C**) The reference model but not the absolute model was also able to predict specific pair-based preferences, which could be generalized across the 3 experiments (3 experiment average) for both learning and transfer phases. This was particularly striking for the critical B|C comparison (red box) during the transfer phase.

**Table 1.**
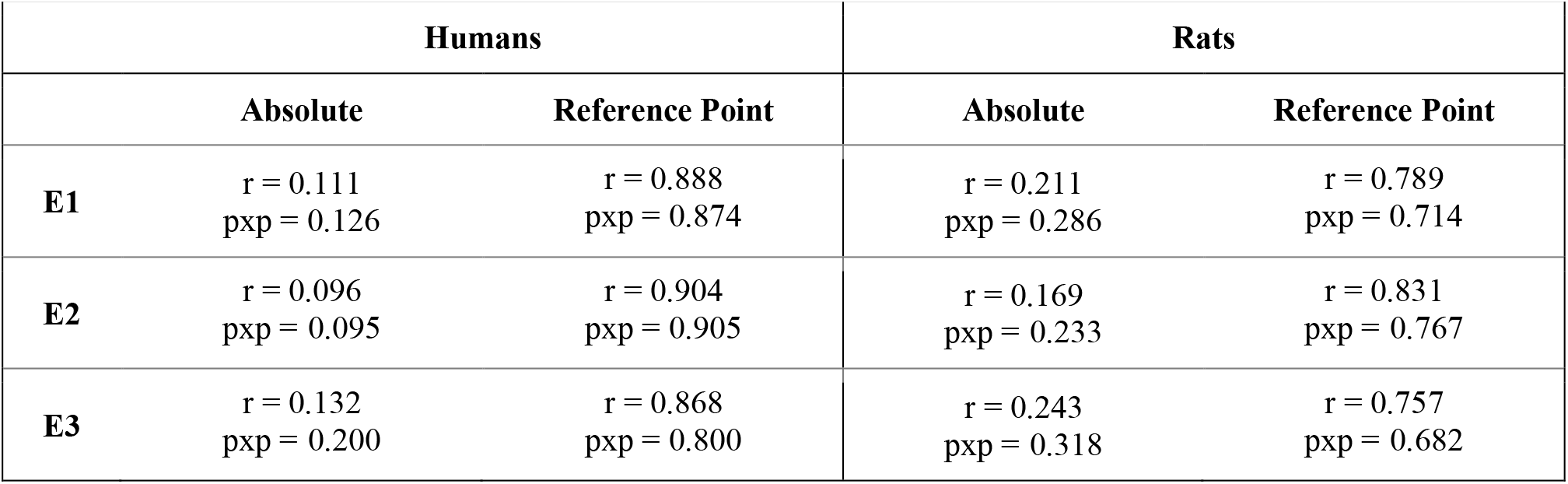
Model Comparison. R refers to the percentage of the sample best described by the given model, while pxp is the probability that the given model best describes the data.

**Table 2.**
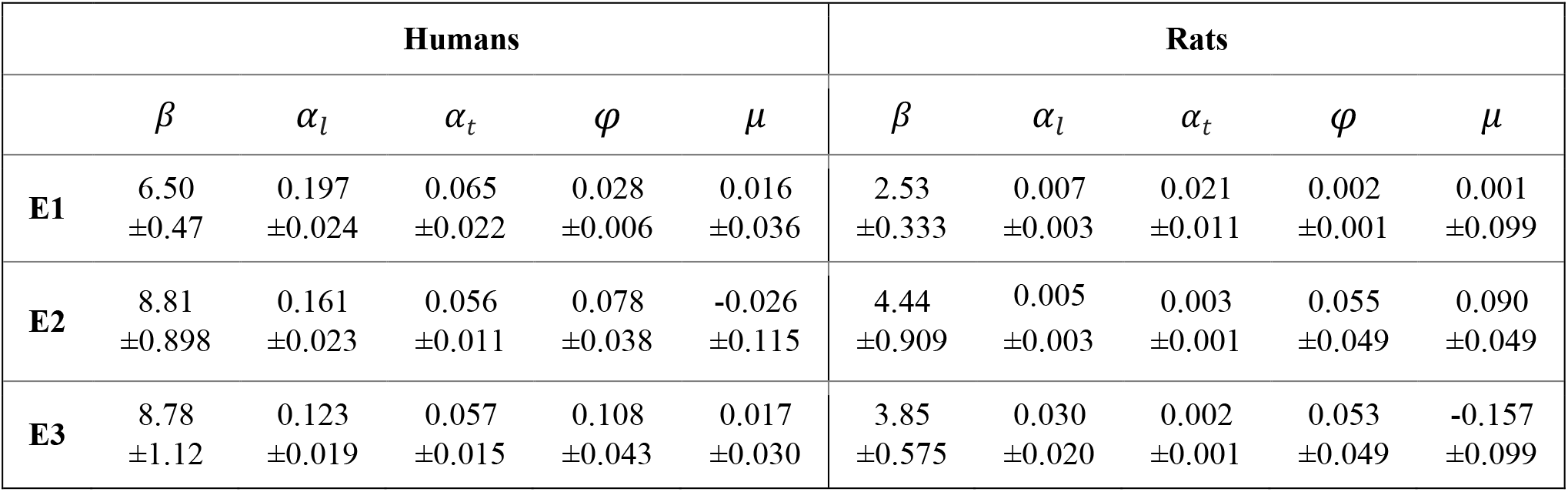
Model Parameters (Mean ± SEM). *β* is the inverse temperature; *α*_a_ is the learning rate of the learning phase; *α*_*t*_ is the learning rate of the transfer phase; *φ* is the forgetting rate; μ is the motor bias.

| Humans |  |  |  |  |  | Rats |  |  |  |  |
| --- | --- | --- | --- | --- | --- | --- | --- | --- | --- | --- |
| | $\beta$ | $\alpha_l$ | $\alpha_t$ | $\varphi$ | $\mu$ | $\beta$ | $\alpha_l$ | $\alpha_t$ | $\varphi$ | $\mu$ |
| **E1** | 6.50<br>$\pm 0.47$ | 0.197<br>$\pm 0.024$ | 0.065<br>$\pm 0.022$ | 0.028<br>$\pm 0.006$ | 0.016<br>$\pm 0.036$ | 2.53<br>$\pm 0.333$ | 0.007<br>$\pm 0.003$ | 0.021<br>$\pm 0.011$ | 0.002<br>$\pm 0.001$ | 0.001<br>$\pm 0.099$ |
| **E2** | 8.81<br>$\pm 0.898$ | 0.161<br>$\pm 0.023$ | 0.056<br>$\pm 0.011$ | 0.078<br>$\pm 0.038$ | -0.026<br>$\pm 0.115$ | 4.44<br>$\pm 0.909$ | 0.005<br>$\pm 0.003$ | 0.003<br>$\pm 0.001$ | 0.055<br>$\pm 0.049$ | 0.090<br>$\pm 0.049$ |
| **E3** | 8.78<br>$\pm 1.12$ | 0.123<br>$\pm 0.019$ | 0.057<br>$\pm 0.015$ | 0.108<br>$\pm 0.043$ | 0.017<br>$\pm 0.030$ | 3.85<br>$\pm 0.575$ | 0.030<br>$\pm 0.020$ | 0.002<br>$\pm 0.001$ | 0.053<br>$\pm 0.049$ | -0.157<br>$\pm 0.099$ |

## Discussion

Context-dependent evaluation appears to be a general feature of decision-making that exists in diverse range of species, including humans (Anlló et al., 2024; Bavard et al., 2018, 2021; Klein et al., 2017; Palminteri et al., 2015), macaques (Lak et al., 2016; Padoa-Schioppa & Assad, 2006; Tobler et al., 2005; Tremblay & Schultz, 1999), rodents (Kuwabara et al., 2020), bats (Hemingway et al., 2021), birds (Bateson et al., 2003; Belke, 1992; Freidin & Kacelnik, 2011; Gibbon, 1995; Herrnstein, 1961; Monteiro et al., 2013; Pompilio & Kacelnik, 2010; Vasconcelos et al., 2013) and insects (Shafir et al., 2002; Solvi et al., 2022; Villar et al., 2022). However, there has been a methodological divide between studies in human and non-human species. First, most non-human studies have focused on the relative distribution of responses to options in the context as function of their reinforcement (i.e., matching law) (Herrnstein, 1961), or to relative neurobiological signals that scale with outcomes (Louie et al., 2015), rather than on direct tests of biases that can emerge following the transfer of learned relative values to new contexts, as is common in the human studies. Moreover, reference-point centering has emerged as a very plausible candidate to explain many context-dependent learning phenomena (Bavard et al., 2018, 2021; Hayes & Wedell, 2023; Juechems & Summerfield, 2019; Palminteri & Lebreton, 2021) in humans, yet it is unknown if the same mechanisms apply in non-human species.

The present work sought to provide direct behavioural and computational cross-species comparisons of context-dependent valuation. In order to achieve this, we designed a two-arm bandit reinforcement learning task that was structurally similar for humans and rats, inspired by previous studies (e.g., Pompilio & Kacelnik, 2010; Bavard et al., 2018). In the learning phase, options were presented in fixed pairs, which defined stable choice contexts, and both humans and rats had to choose the option associated with the higher reward probability. They then performed a transfer test during which all possible binary combinations were presented, including novel choice contexts. Again, humans and rats were required to select the option with the higher reward probability. Both species successfully chose the high value options during the learning phase; however, this preference emerged more quickly in humans than in rats. Choices during the subsequent transfer test did not, however, follow the absolute values. When presented with two options associated with the same reward probability, both species chose the option that had previously been paired with a lower value option. In addition, both species exhibited suboptimal preferences such that they preferred an option with low absolute value but high relative value over an option with high absolute but low relative value. Both human and rat subjects corrected these relative value biases over subsequent transfer trials but at different rates, with humans showing a faster bias-correction rate than rats. Thus, using the same task in humans and rats, we demonstrate that relative value learning can represent a predictable source of systematic suboptimal decisions; i.e., choices that do not maximize the expected future rewards, and that rates of recovery from these biases diverge in humans and rats.

Our results contribute to the extensive evidence that contextual effects are pervasive in learning. This suggests that such effects represent an adaptive response shaped by shared neurobehavioural adaptations to similar environmental and physiological constraints across species. For instance, evolutionary simulations have demonstrated that relative memories of outcomes may be adaptive in environments that alternate between periods of high and low average rewards (e.g., “spring” versus “winter”) (McNamara & Houston, 2009). In rich environments, sharp memory of positive outcomes is less crucial, whereas in poor environments, remembering the few positive rewards with heightened (relative) value becomes essential for survival. Hunter and Daw (2021) also proposed that relative, reference point-dependent valuation offers ecological advantages by approximating the opportunity cost of choices in foraging situations where multiple options exist but cannot be explored fully (Hunter & Daw, 2021). The similarities that we observed in humans and rats, that occupy very different ecological niches, suggests that the environmental or neurobiological constraints shaping reference-point centering are sufficiently common and shared across species.

At the neural level, reference-point centering can also be understood as a form of normalization, underpinning an adaptive mechanism for coding external signals. Such normalisation has been thought of as a fundamental neurocomputational mechanism for efficient information coding in the brain (Burke et al., 2016; Louie et al., 2013; Louie & Glimcher, 2012; Padoa-Schioppa & Rustichini, 2014; Rustichini et al., 2023). In this way, normalisation via reference-point centering may reflect a homeostatic setpoint of neural responses to the available options within a context, tuned by reinforcement learning processes (Juechems & Summerfield, 2019; Keramati & Gutkin, 2014). Moreover, neural systems face well-known constraints, such as refractory periods and the inability to encode negative firing rates, which limit the efficiency of encoding information such as prediction errors (Bayer & Glimcher, 2005; Palminteri & Pessiglione, 2017). Rescaling outcome encoding relative to a reference point addresses these limitations by adjusting to the average value in a manner analogous to how the pupil adapts its sensitivity to average luminosity. Thus, our findings highlight how reference-point dependent valuation reflects a convergence of ecological and neural adaptations that enhance learning efficiency across species.

Our results demonstrate that the same computational process, reference-point centering—originally developed to explain behavioural and neural rescaling effects across gains and losses in humans—can be effectively adapted to account for normalization effects in tasks featuring varying probabilities of positive outcomes across species (Palminteri et al., 2015). The preservation of this computational process, despite some inevitable adaptations in the task structure (such as an increased number of trials and the shift from secondary to primary rewards), underscores its robustness. Furthermore, this process is not confined to reinforcement learning; appearing in other domains, such as perceptual judgment (e.g., adaptation level theory by Helson) and decision-making under risk, where reference point dependence is a central tenet of prospect theory (Helson, 1948, 1964; Kahneman & Tversky, 1979).

The ubiquity of reference point centering across these diverse domains further supports the idea that it represents a canonical neural computation (Carandini & Heeger, 2012; Chirimuuta, 2014). Additionally, our findings contribute to the ongoing debate about the functional form of value normalization by demonstrating a high degree of robustness to reference point dependence across species. This robustness aligns with findings from other laboratories using different tasks, reinforcing the generality and importance of this computational mechanism (Hayes & Wedell, 2023). Beyond value normalization, in both species, we observed that learning rates decreased when moving from the learning to the transfer phase. This is reminiscent of the machine learning technique of “annealing”, where learning rate reduction over training is shown to display adaptive value (Cagan & Kotovsky, 2002; Zeiler, 2012). This is also consistent with the idea that both species naturally appraise and parse their experience with the task in an initial phase of rapid belief formation and the later phase of slow revision of these beliefs.

Despite many similarities, we did observe some quantitative behavioral differences between the two species. First, rats appeared more sensitive to actual reward probabilities for the bad options compared to the good options, particularly when deterministic outcomes were used. While humans showed similar learning performance in the rich [AB] and the poor [CD] contexts, rats showed better learning in the poor context. Moreover, during the transfer trials, humans tended to be more accurate when choosing between the previously good options [AC]. Conversely, this was not the case for rats, who displayed higher accuracy when choosing between the previously bad options [BD]. This suggests that rats ‘recall was better in the “poor” environment, compared to the “rich” environment, which perhaps reflects a greater frustration (loss aversion) in rats than humans, likely due to the non-delivery of expected primary rewards in rat experiments (Amsel, 1962; Papini et al., 2022).

Second, humans overcame their context-dependent biases across the transfer phase by re-ranking the options according to the new context and their absolute value. By contrast, rats tended to correct their bias only when not doing so led to suboptimal performance. That is, rats ‘preference for C over B diminished when C had an objectively lower reward value but not when B and C were associated with equal reward probabilities. This is well-explained and captured by the difference in learning rates between the two species. However, it is possible that this persistence reflects the incipit of habit formation, especially given the longer duration of training in rats compared to humans (Miller et al., 2019; Pompilio & Kacelnik, 2010; Sugawara & Katahira, 2021). Nevertheless, it must be noted that, in the case of equal reward probabilities, there is no immediate cost associated with maintaining a preference for the option that had higher relative value, and rats may have eventually overcome this bias with extended training in the transfer phase.

Direct comparisons of model parameters are complicated by some necessary structural differences in the tasks—most notably, during the learning phase, where rats experienced forced choices and partial feedback, while humans had free choices and complete feedback. Nevertheless, we did observe substantially different parameter values across the two species. Specifically, humans exhibited higher learning rates and choice temperatures compared to rats. Since higher learning rates are particularly associated with working memory activity and representations (Collins & Frank, 2012), whereas lower learning rates are linked to striatal processes and slower associative learning (Balleine et al., 2007; Zwosta et al., 2018), we speculate that these differences may be partly explained by a greater involvement of the prefrontal cortex in humans compared to rats.

In our task, options are compared along a single dimension (i.e., probability of reward). It remains to be determined if the choice behaviour of humans and rats is as similar if other outcome dimensions are manipulated, including reward magnitude or delay and whether or not the same process of normalization—reference point dependence—will apply these scenarios (Bavard & Palminteri, 2023; Louie et al., 2015). To ensure that given equal information about option values was provided, we used a full feedback (i.e., choice-independent) procedure in humans and forced sampling trials in rats. We measured rats ‘preferences in the learning phase by including unrewarded probe choice trials. Given that both species made choices during the learning phase, it remains possible that they formed preference habits (Miller et al., 2019; Pompilio & Kacelnik, 2010; Zwosta et al., 2018). That is, options may have been preferred at transfer simply because they had been chosen more often during the learning phase. However, in rats, these probe choice trials were unrewarded and thus, if anything, this reduced the experienced reward probability of C versus B (as C was chosen more often that B in the probe trials during learning). Furthermore, given the short timeframe of the task, habit formation is unlikely to explain the observed effects in humans, as relative valuation emerged within just a few minutes (de Wit et al., 2018). Thus, attributing the same behavioural phenomena (e.g., preference for C in BC) to two distinct mechanisms—habit formation in rats and relative valuation in humans— appears far less parsimonious than considering the results in both species as arising from the same underlying computational cognitive process.

Having demonstrated similar behavioural biases and computational processes in rats and humans, our findings pave the way for significant future translational endeavors. First, while there is extensive neurobiological evidence for the relative encoding of option values across various species (including humans, macaques, and rodents) in regions such as the dopaminergic midbrain, orbitofrontal cortex, striatum, and amygdala (Gardner et al., 2019; Lak et al., 2016; Morris et al., 2006; Padoa-Schioppa & Assad, 2008; Rangel & Clithero, 2012; Syed et al., 2016; Tobler et al., 2005; Tremblay & Schultz, 1999; Xie & Padoa-Schioppa, 2016), the neural mechanisms underlying the encoding of the reference point itself remain much less understood and underexplored. Furthermore, recent studies show that context-dependent reinforcement learning is impaired in certain pathologies, such as addiction (Gueguen et al., 2024). By successfully replicating these findings in rats, we propose that our task and computational model could serve as a valuable behavioural marker for translational research aimed at improving our understanding of these clinical conditions.

Overall, our study illustrates the feasibility of direct computational comparative studies, highlighting their potential for deepening our understanding of conserved and robust cognitive processes. The approach used in this study of minimising task and reinforcement learning structure differences between species holds significant translational potential, offering valuable insights for both neuroscientific and clinical research.

## Methods

### Subjects

#### Human Participants

Participants were recruited via the online platform Prolific. In total, 72 participants completed the experiment, i.e., 24 per experiment (36 females, age: M=29.9, SD=10.6). Participants were paid £2.50 plus a bonus payment up to £2.34 (M=1.98, SD=0.181) based on the number of points they won during the experiment. See **Supplemental Table 4** for mean age, sex, completion time and bonus payment. The study was approved by a local ethical committee and all participants gave informed consent before taking part in the experiment. We excluded two human participants from the model fitting due to missing data.

#### Animals

A total of 44 Long-Evans rats (36 males and 8 females) aged 3-4 months were acquired from Janvier, France. The rats were pair-housed and allowed two weeks to acclimate to the laboratory environment before experimentation began. The facility maintained a steady temperature of 22±1”C with a 12-hour light/dark cycle (lights on from 07:00-19:00). The rats underwent daily handling for five days before behavioural tasks and were placed on food restriction two days prior to and throughout the behavioural procedure. Every 24 h, following behavioural training, approximately 12 grams of standard lab chow per animal was left in their home cage. Rats were weighed and monitored to ensure they maintained

∼90% of their free-feeding body weight and a healthy disposition. The experimental procedures were performed in compliance with both French (council directive 2013-118, dated 1st February 2013) and international (directive 2010-63, dated 22nd September 2010, European Community) legislations.

### Experimental design

#### Apparatus

Human experiments were conducted online using a phone, tablet, or a computer. Rats were trained in 4 identical operant cages (40 x 30 x 35 cm; Imetronic, France), each situated within a light and sound-attenuating wooden enclosure (74 x 46 x 50 cm). Each cage consisted of two opaque panels on the left and right, two transparent Perspex walls on the front and back, and a stainless-steel grid floor (rod diameter: 0.5 cm; inter-rod distance: 1.5 cm). Cages were fitted with a pellet dispenser that, when activated, delivered sucrose pellets (45 mg, Test Diet, catalog # 1811251) into a central food port (6 x 4.5 x 4.5 cm) on the right wall. A retractable lever (4 x 12 cm) was also fitted to the left of the food port. A touchscreen (11.6 inch, LCD display, 1920*1080 resolution, capacitive touch type; Waveshare Electronics) was situated on the left wall, opposite the food port and lever. The touchscreen display was fitted horizontally at the base of the wall. All stimuli (10 x 10 cm) were presented on the centre-left or centre-right of the screen. Data acquisition and equipment control were managed by POLY software on a computer connected to the operant chambers (Imetronic, France). Touchscreen data was collected on a trial-by-trial basis, including the correct and incorrect responses, omitted trials, and response latencies.

#### Stimuli

Human experiments used 8 visual stimuli, while only 4 of these stimuli were used in the rat experiments. In human experiments, the stimuli were assigned to options at random for each participant. Likewise, in rats, the stimuli assigned to each option were counterbalanced such that each stimulus could be option A, B, C, or D. To enhance the discriminability of the stimuli while simultaneously mitigating any innate preferences, stimuli were created with varying features held at equidistant degrees of difference. Given the capacity of rats to detect colours within their visual spectrum and to discriminate according to line orientation (Jacobs et al., 2001; Rocha et al., 2016; Wiesbrock et al., 2022), we chose colored Gabor patches. We defined 3 stimulus feature characteristics: Gabor filter orientation, frequency, and background color (**Supplemental Table 5**). Other characteristics were held common, including size (pixels = 100), envelope (Gaussian), sigma (σ = 30), phase (0), and pattern colour (colour 1 RGB = 191.25; colour 2 RGB = 63.75) The stimuli were generated using the open access Gabor patch generator hosted at (Mathot, 2014/2025): http://www.cogsci.nl/software/online-gabor-patch-generator.

### Behavioural procedure: Humans

#### Learning Phase

Human participants were presented with two options on the screen and were required to select one of them on each trial. Crucially, after each choice, participants received full feedback, i.e., outcomes of both the chosen and unchosen options were displayed on the screen. However, only the outcome of the chosen option was added to the participant’s score. This design allowed us to reduce the number of trials while ensuring that participants have the same amount of information about each option, as was the case in the rat learning phase.

The reward probabilities associated with each option are shown in **Figure 1C**. Two stimuli were associated with each probability (8 options in total) to ensure that the task remained sufficiently challenging. The side on which the options were presented was pseudo-randomised (equal number of presentations on each side) as was the order of the outcomes. Each option was shown 32 times in four blocks of 8 trials, yielding 128 trials in total. To create the AB versus CD contexts, participants were presented with block-wise presentations of either AB or CD choices over a single session.

To maximise participants ‘performance, participants were told at the start of the experiment that they would encounter the same options in all phases, and that any knowledge gained in the learning phase will be useful in the subsequent phases of the experiment. However, we did not disclose the exact nature of the subsequent phases. Furthermore, the learning phase was preceded by a pre-training phase consisting of 16 trials during which the participants could familiarize themselves with the task structure. The pre-training phase used a different set of stimuli (the letters A, B, C, D) with different outcome probabilities (25% and 75%).

#### Transfer Phase

During the transfer phase, participants were presented with the same options as in the learning phase, however, the options were shown in all possible pairs within the cluster; i.e. A was presented with B, C or D but not with A ‘or B’. Participants were asked to select one of the two options presented on the screen and they received only partial feedback (i.e., the outcome of the selected option). There were 120 trials in total.

#### Explicit Recollection Task

The probability recollection task, specific to the human participants, was designed to assess to what extent participants remembered the exact reward probability or frequency of each option. Each option was presented three times in a pseudorandom manner, and participants were required to estimate its reward frequency using a slider scale from 0 to 100% (See **Supplemental Materials** for more details).

### Behavioural procedure: Rats

#### Pre-training

Rats were first trained to retrieve sugar food pellet rewards from the food port. In these sessions, rats were placed in their allocated operant cage and 10 sugar pellets (45 mg, 3.35 kcal/g each) were delivered to the food port every 60 s. Rats underwent one food port training session per day for three days.

On day 4, rats began their touchscreen training. During sessions 1-4, a single stimulus (a white square) was randomly presented on either side of the touchscreen screen, and a reward was delivered when the rat made physical contact (i.e., touch) with the white square stimulus. Touches made on the screen that were not targeted towards the stimulus were not rewarded. Each of these sessions included 10 trials with a maximum duration of 30 minutes. Rats received one session per day for 4 consecutive days. In sessions 5 and 6, the touchscreen was turned off and a lever, located to the right of the food port, was extended and rats were required to press the lever to earn the sugar pellet reward. Finally, in sessions 7-12, rats were required to integrate their learning from the previous sessions and were trained to press the lever to initiate the stimulus presentation on the touchscreen. During these final sessions, a single press on the lever now initiated the stimulus presentation and a single touch on the white square stimulus presented on the touchscreen (side-randomised) earned the reward. Each session in this phase consisted of 30 trials with a maximum duration of 45 minutes.

#### Learning Phase

After pre-training, rats were then trained to learn the reward probabilities associated with 4 distinct touchscreen stimuli: A, B, C, D. Rats received two separate sessions per day for 20 days: an AB session and a CD session. One session was conducted in the morning (e.g., the AB session) and the other was conducted in the afternoon (the CD session). The order of the sessions was counterbalanced across days. Each of these sessions involved 96 forced sampling trials and 12 unrewarded probe choice trials. Rats were required to press on the initiation lever to begin each sampling and choice trial. Rewards were delivered during the sampling trials based on the probabilities shown in **Figure 1C**. Choice trials were unrewarded.

Each of learning phase sessions were arranged into 12 blocks. Within each block, 8 sampling trials (single stimulus presentations) preceded a probe (unrewarded) choice trial. Sampling trials included 4 consecutive presentations of the high value stimulus (A or C) and 4 consecutive presentations of the low value (B or D) stimulus, order counterbalanced. The location of each stimulus on the touchscreen (left or right) was fully counterbalanced as was the order of stimulus presentation. That is, each stimulus was presented an equal number of times on the left and right of the screen (during both sampling and choice trials) and we alternated both within and between sessions whether the high value (A or C) or low value (B or D) stimulus was presented first in the sampling trials.

#### Transfer Phase

In the transfer phase, rats were offered choices between the six possible combinations of stimuli pairs, including the intra-context choices (AB, CD) and inter-context choices (four novel pairs = AC, AD, BC, BD). Stimuli were presented in blocks of 12 trials, such that each possible pair was presented twice with the side of the stimuli counterbalanced. Each session included a total of 8 blocks (i.e., 96 choice pairs in total) and the order of the six pairs within each block was pseudorandomised. As for the training phase, rats initiated each trial by pressing on the lever and their choice between the two stimuli in each pair was rewarded based on the probabilities associated with the chosen stimulus.

### Analyses

#### Behavioural Analyses

To ascertain that humans and rats learned to select the correct option in the learning phase, we fitted a generalized linear mixed model (GLMM) with correct response (resp_cor) as the dependent variable, and experiment (exp), choice pair (ch_pair) and choice trial (humans, log-transformed) or session (rats) and their interactions as independent variables. The model also contained a random intercept and an individual slope for the trial/session for each subject (partID). An identical analysis was also performed for the transfer phase.

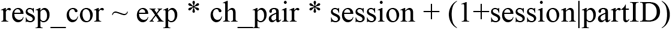

We used the estimated marginal means to determine whether responses were above chance level in each choice pair and whether there were any significant differences between them. In particular, we were interested in the responses in the BC pair, as a strong preference for C would indicate relative encoding, while preference for B (E2) or indifference (E1, E3) would suggest absolute encoding. We examined both average performance across all transfer phase trials as well as performance during the initial transfer trials before any new learning could occur. In all cases, models were fitted separately to the human and rat data. When appropriate, p-values were adjusted using the multivariate t-distribution method.

Furthermore, we also analysed the reaction times in both rats and humans to provide further insight into the underlying cognitive processes (See **Supplemental Materials** for more details).

### Computational Models

We analysed the data with several different models, which mainly differed in how the choice outcomes were encoded. The absolute model assumes that choice outcomes are evaluated independently of other outcomes (absolute encoding), and reference point model assumes that the outcome value is adjusted as a function of the other outcomes presented at the same time or in the near past (relative encoding).

### Absolute Learning Model

The Absolute Learning Model and all undermentioned models are variants of a Q-learning model (Palminteri & Lebreton, 2021; Sutton & Barto, 1998) which assumes that subjects calculate absolute values (Q-values) of all options based on their trial-by-trial outcome history and then use these Q-values to decide which option to pick next. The absolute model in particular, assumes that outcomes are encoded independently of other values, i.e. exactly as seen.

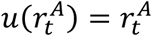

The value update can be described as:

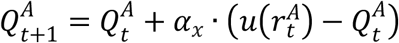

Where 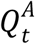 is the estimated value of option *A* at a trial 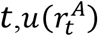 is the subjective outcome of the option, which in this case equals to the objective outcome, and *α*_*x*_ is the learning rate that can take two values as function of the phase (learning: *α*_*l*_; transfer:*α*_*t*_). We fitted two learning rates in this model, one was utilised in the learning phase and one in the transfer phase to account for potential learning differences between the phases.

To account for the fact that subjects might forget the value of an option if they have not seen it for a while, we added a decay parameter *φ* to the model that reduced the Q-values of unseen options towards 0, which represented a mid-point in our fitted data.

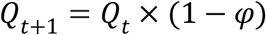

To model the decision phase, we used a softmax decision rule which combined the value of an option with a selection bias for the side it was presented on.

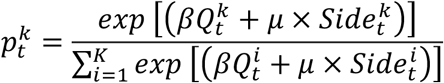

Where 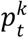 is a probability to select option *k* at trial 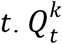 is the estimated value of option *k* at trial *t* and *β* is the inverse temperature, a parameter that regulates the preference for the option with the highest estimated value. At *β* → ∞, subjects would always choose the option with the highest absolute value, whereas at *β* = 0 subjects would treat all options equally and choose at random. In addition, the model also includes a motor bias parameter μ, which is the bias for selecting options on the left. Finally, 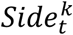 refers to the side option *k* was presesented on at trial *t* (left ∼ 1, right ∼ -1).

#### Reference Point Learning Model

The outcomes in the reference point model are scaled as a function of the average outcome in the given context (i.e. choice pair). The relative outcome is calculated in two steps. First, the reference point is updated, then the updated reference point is used to scale the outcomes.

If subjects receive full feedback, i.e. outcome for both the chosen and unchosen option is shown, the reference point is calculated as follows:

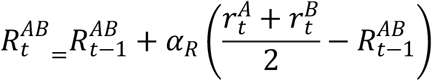

Where 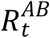 is the reference point for context AB at 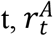 and 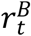 are outcomes of option A and B at trial t, and *α*_*R*_ is the reference learning rate, which governs how fast is the reference point updated. In our study, the value of the *α*_*R*_ parameter was fixed to a mean of the learning rates for the learning and transfer phases.

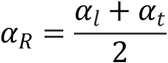

If subjects received only partial feedback the reference point was based on the outcome of the chosen value and the last outcome seen for the unchosen value (outcome from when the option was last chosen).

Once the reference point is calculated, the reference is subtracted from the real outcome.

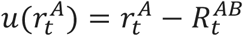

The subjective outcome, *u*(*r*_*t*_), is then used to update the Q-values which are subsequently used to make a decision. This process is identical in both absolute and reference point models. Note, that in the reference point model both the Q-value and remembered value of the last seen value outcome decayed at the same rate.

#### Parameter optimisation and model comparison

To estimate parameters values, we optimized the maximum log likelihood using the Nelder-Mead algorithm. We used weakly informative priors for each parameter to avoid degenerate parameter estimates, and five starting points to increase the probability of finding the global optimum.

The parameter priors were based on previous studies and were set to:

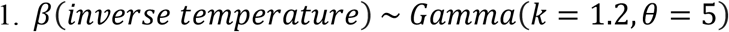

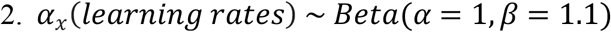

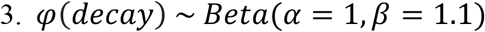

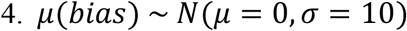

To simplify the optimization process, we transformed the outcomes from the (0,1) range to the (-1,1). This allowed us to initialize the Q-values at 0 for all models. In addition, we assumed that, with the exception of the learning rates, parameter values would be same for the learning phase and the transfer phase.

Once we obtained the best set of parameters, we calculated Schwarz (BIC) weights for each model and subject and compared them using Bayesian model selection. This allowed us to estimate protected exceedance probability (pxp) of each model, i.e. probability, corrected for chance, that a given model describes the data better than any other model in the set. In addition, we validated the models by comparing their predictions against the actual data by simulating each subject using their best-fitted parameters and examining to what extend the simulated data can reproduce the key behavioural findings.

#### Parameter and model recovery

The parameter recovery was performed by simulating data of 100 synthetic subjects based on the reference point model. The simulated data were then refitted using the same model and fitting procedure. For model recovery, we simulated data of 100 synthetic subjects based on each of the models. Each of these generated datasets was then fitted by all other models, and the best fitting model for each subject was determined by BIC. We then calculated the confusion matrix, i.e. the probability that a model fits best given the true generative model, and inversion matrix, i.e. the probability that the data was generated by a specific model, given it is the best fitted model. The model and parameter recovery was satisfactory for both humans (**Supplemental Figure 4**) and rats (**Supplemental Figure 5**).

### Software

Statistical analyses were performed using R (version 4.1.3) and the following packages: afex (Singmann et al., 2022), DHARMa (Hartig, 2022), emmeans (Lenth et al., 2022), glmmTMB (Brooks et al., 2017) and performance (Lüdecke et al., 2021), for behavioural analyses. For fitting the learning models we used optimx (Nash & Varadhan, 2011) to find the best parameter values, and bmsR (Lisi, 2021) to find the best model. All graphs were plotted using ggplot2, which is part of a wider tidyverse package collection (Wickham et al., 2019) that was used for general data manipulation. Additionally, we used parallel (R Core Team, 2022) and future (Bengtsson, 2021) to enable parallel processing during the fitting of the learning models.

## Supporting information

Suppelmentary Information

## Acknowledgements

Shauna Parkes thanks Yoan Salafranque for animal care and Niniva Ghosh for assistance with rat behavioural experiments. Stefano Palminteri acknowledges support from the European Research Council consolidator grant (RaReMem: 101043804) and the Agence Nationale de la Recherche grants (RANGE: ANR-21-CE28-0024-01). Shauna Parkes and Stefano Palminteri acknowledges support from Agence Nationale de la Recherche grants (RELATIVE: ANR-21-CE37-750 0008-01).

