## Supplementary material for "Reference Point-Dependent Reinforcement Learning in Humans and Rats": Suppelmentary Information

### Supplemental Materials

#### Response times

We analysed subjects' reaction times as such analysis can provide further insight into the underlying cognitive processes. Notably, longer reaction times are linked with decision conflict or option values (Fontanesi et al., 2019). After assessing that we could observe the expected modulation of reaction times as a function of the progression in the learning phase (**Figure S6**), we focussed on analysing the reaction times as a function of the option pair, after averaging across trials.

To do so, we first excluded any trials with response times under 100ms as the former were too short to represent a genuine response to the presented stimuli. We also excluded reaction times over 5s (humans) or 10s (rats) as those were likely result of task-unrelated events. The remaining trials were analysed using the following linear mixed model (LMM):

log(respRT) ~ expID*context+context*blockTrial+context*repetition+(1|partID)

Where respRT was the response time in ms, context was the presented pair of options, e.g. AB or CD, blockTrial represented a trial within the miniblock (humans) or session (rats), and repetition referred to the session (rats) or to the miniblock (humans). Both blockTrial and repetition variables were centred. Note that blockTrial was used for the learning phase only (as the transfer phase had no blocked content), and the interaction between context and repetition was omitted in the human transfer phase due to collinearity issues.

In the learning phase (**Table S6**), rats had significantly longer reaction times in the "poor" (CD) context, compared to the "rich" (AB) context even though the objective choice difficulty, proxied by the probability difference between the presented options, was the same in both contexts (E1: $contrast_{CD/AB} = 1.12$, $t(20787)= 12.9$, $p< 0.001$; E2: $contrast_{CD/AB} = 1.17$, $t(20787)= 20.2$, $p< 0.001$; E3: $contrast_{CD/AB} = 1.07$, $t(20787)= 9.18$, $p< 0.001$). However, this was not the case for humans (E1: $contrast_{CD/AB} = 0.968$, $t(9072)= -2.18$, $p= 0.02$; E2: $contrast_{CD/AB} = 0.997$, $t(9072)= -0.204$, $p=.839$; E3: $contrast_{CD/AB} = 0.959$, $t(9072)= -2.78$, $p= .005$). One possible explanation is that rats viewed the poor contexts as more punishing than humans as response times are known to be slower in such conditions (Fontanesi et al., 2019; Palminteri et al., 2016).

In the transfer phase, both species responded slowest in the BD context, where they had to decide between two previously unfavourable options (**Table S7**). This is notable as the objective difficulty (again proxied by the probability difference between the options) was the same as in the AC context, where subjects encountered two previously advantageous options and where the response times were significantly faster (**Humans**: E1: $contrast_{BD/AC} = 1.24$, $t(8297)= 7.5$, $p< 0.001$; E2: $contrast_{BD/AC} = 1.21$, $t\ (8297)=6.44$, $p< 0.001$; E3: $contrast_{BD/AC} = 1.11$, $t(8297)= 3.62$, $p< 0.001$, **Rats:** E1: $contrast_{BD/AC} = 1.04$, $t(42028)= 3.52$, $p< 0.001$; E2: $contrast_{BD/AC} = 1.05$, $t\ (42028)=5.18$, $p< 0.001$; E3: $contrast_{BD/AC} = 1.06$, $t(42028)= 6.46$, $p< 0.001$)

Notably, these results cannot be explained by a difference in information sampling, since both human and rat subjects were provided with complete feedback in the learning phase and these effects are present from the first trial of the transfer phase and confirm that in reinforcement learning, option valence is a strong modulator of reaction times (of note, similar findings were recently reported by Tohidi-Moghaddam & Tsetsos (2024))

#### Explicit ratings

The probability recollection task, specific to the human participants, was designed to assess to what extent participants were able to retrieve the probability of reward associated with each option ("*how frequently the option won*"). Each option was presented three times in a pseudorandom manner, and participants were required to estimate its reward probability using a slider scale from 0 to 100%.

In general, participants had a tendency to assign higher reward probabilities to the best possible option (A) and the lower probability to the worst possible option (D), indicating that participants were able to retrieve sensible numbers. However, interestingly, the intermediate value options B and C were encoded in a way that did not follow their objective values (**Figure S7A**).

To quantify this effect, we analysed to what extent the recalled rating differed from the real reward probability for the mid options B and C using a linear mixed model (LMM) with response error and experiment as the independent variables:

resp_diff~symID*expID+(1|partID)

Symbols A and D were excluded from the analysis as they were the extreme options in experiments E1 and E2 and therefore their error was always one directional.

We found that option B, which was disadvantageous in the original learning context, tended to be remembered significantly worse compared to option C, which was advantageous in the original learning context (E1: contrast_B-C = -0.190, $t(856)=-7.34$, $p<0.001$; E2: contrast_B-C = -0.269, $t(856)=-10.4$, $p<0.001$; E3: contrast_B-C = -0.04, $t(856)=-1.64$, $p=0.101$, **Figure S7B**). This is even more striking if we take into account that the explicit rating task was performed after the transfer phase, giving participants the opportunity to correct possible context-dependent biases.

In addition, a significant proportion of participants' choices in the last round of the transfer phase, was consistent with the average option ratings in the probability recollection task, which followed immediately after (E1: M=74.8%, $t(23)=7.04$, $p<0.001$; E2: 69.8%, $t(23)=3.48$, $p=0.002$; E3:73.6%, $t(23)=5.83$, $p<0.001$; two-sided one-sample t-test all compared to the chance level of 0.5).

We can thus conclude that context in which options were presented affected not only participants' choices but also their recollection of the reward probabilities. These results corroborate previous findings on this topic (Bavard & Palminteri, 2023; Juechems et al., 2022; Soukupova et al., 2024).

#### Supplemental Figures

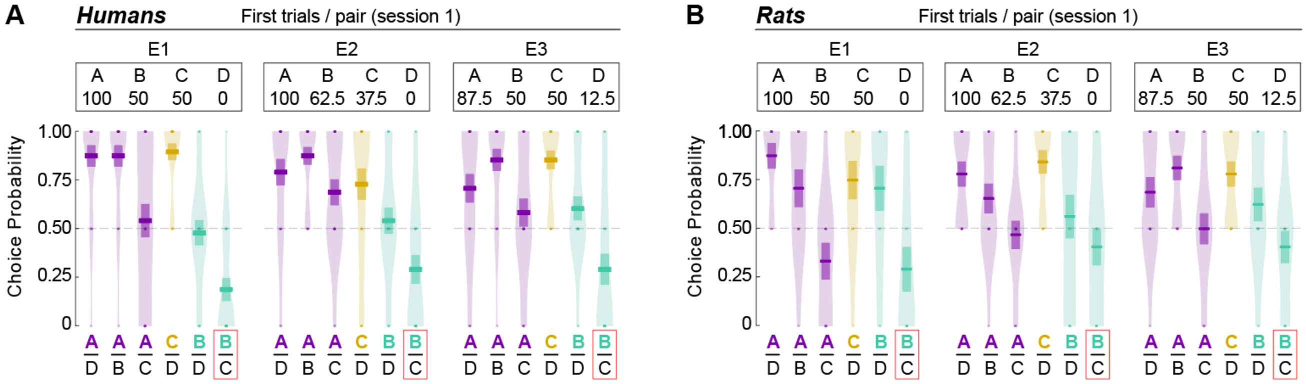


**Supplemental Figure 1**: Initial transfer phase trials for humans and rats. The plots show choice probability for all possible option combinations in the initial round of the transfer phase in humans (**A**) and rats (**B**) for all three experiments (E1, E2, and E3). I.e. choices subjects made when they encountered the given choice pair for the first time in the transfer phase. Individual data points represent the average performance per subject (2 trials/pair). Each pair in the plot is represented by two choices, as humans saw two sets of symbols representing each choice pair (e.g., AD and A'D') and in rats, this corresponds to the counterbalanced side presentation (e.g., one AD trial with A on the left, and one trial with A on the right).

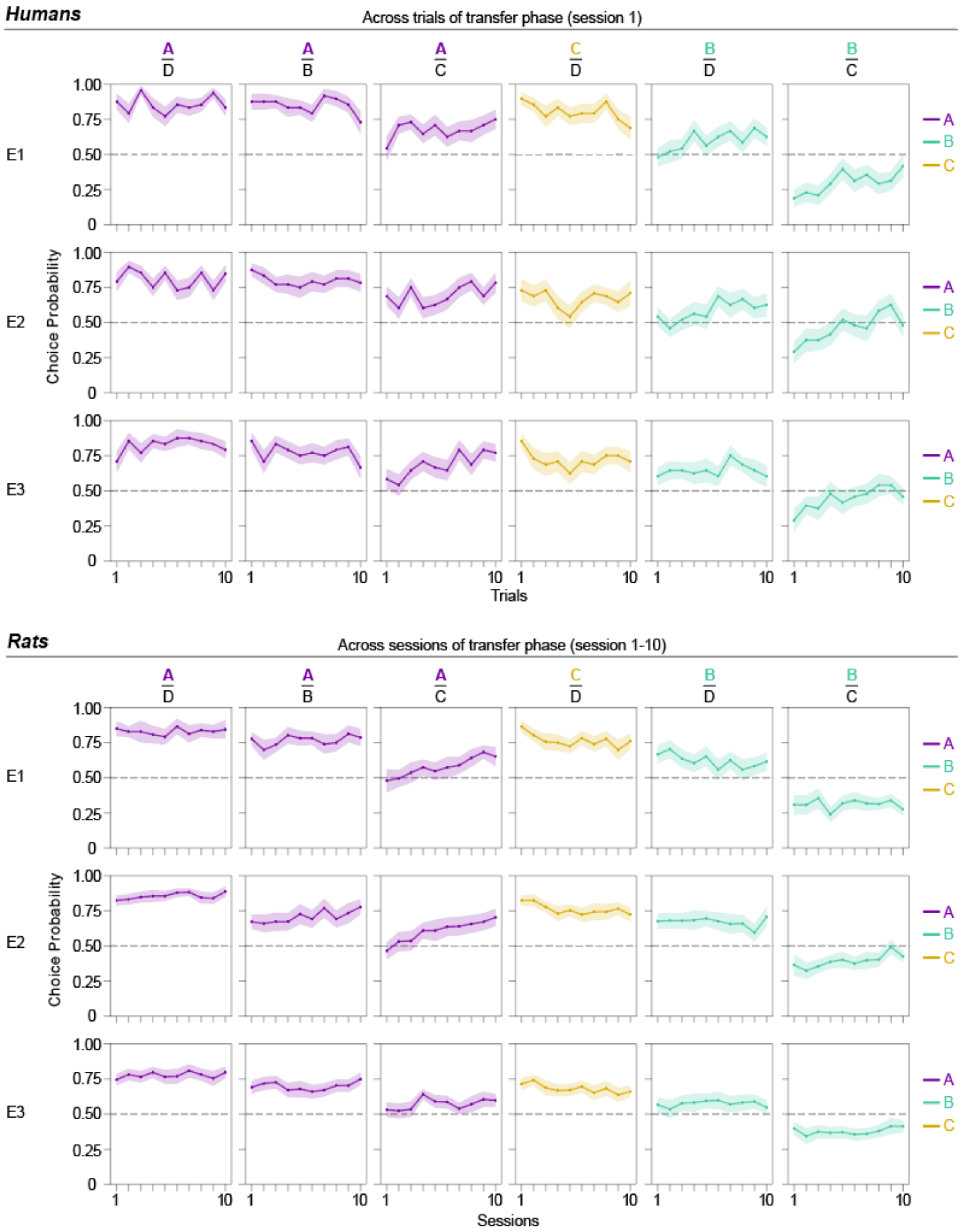


**Supplemental Figure 2**: Evolution of transfer phase choice across all trials (humans) and all sessions (rats). Each point represents the average choice probability for a given trial or session, color represents the option chosen, and the ribbon represents the standard error of the mean. Note that economically suboptimal preferences get corrected over time (e.g., BC – in E2 - and BD in humans, AC and BC in rats).

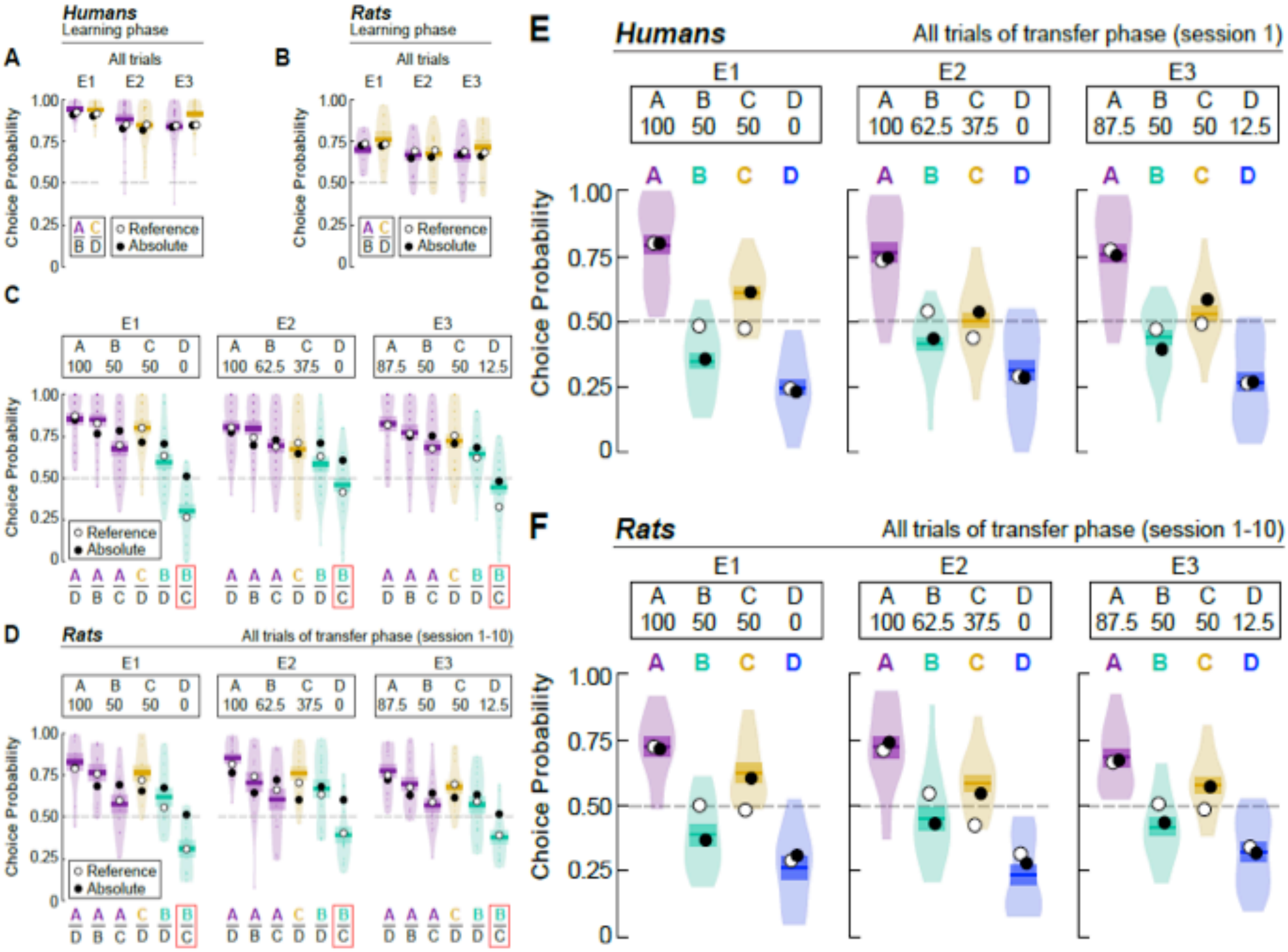


**Supplemental Figure 3**: Behavioral results and model simulations by experiment. Crossbars and violin plots represent subject's responses, while black and white dots represent predictions of the "Absolute" and "Reference" models, respectively. (**A, B**) Choice rates in the learning phase in humans and rats. (**C, D**) Choice rates in each choice pair of the transfer phase in humans and rats. (**E, F**) Choice rate per option in the transfer phase in humans and rats.

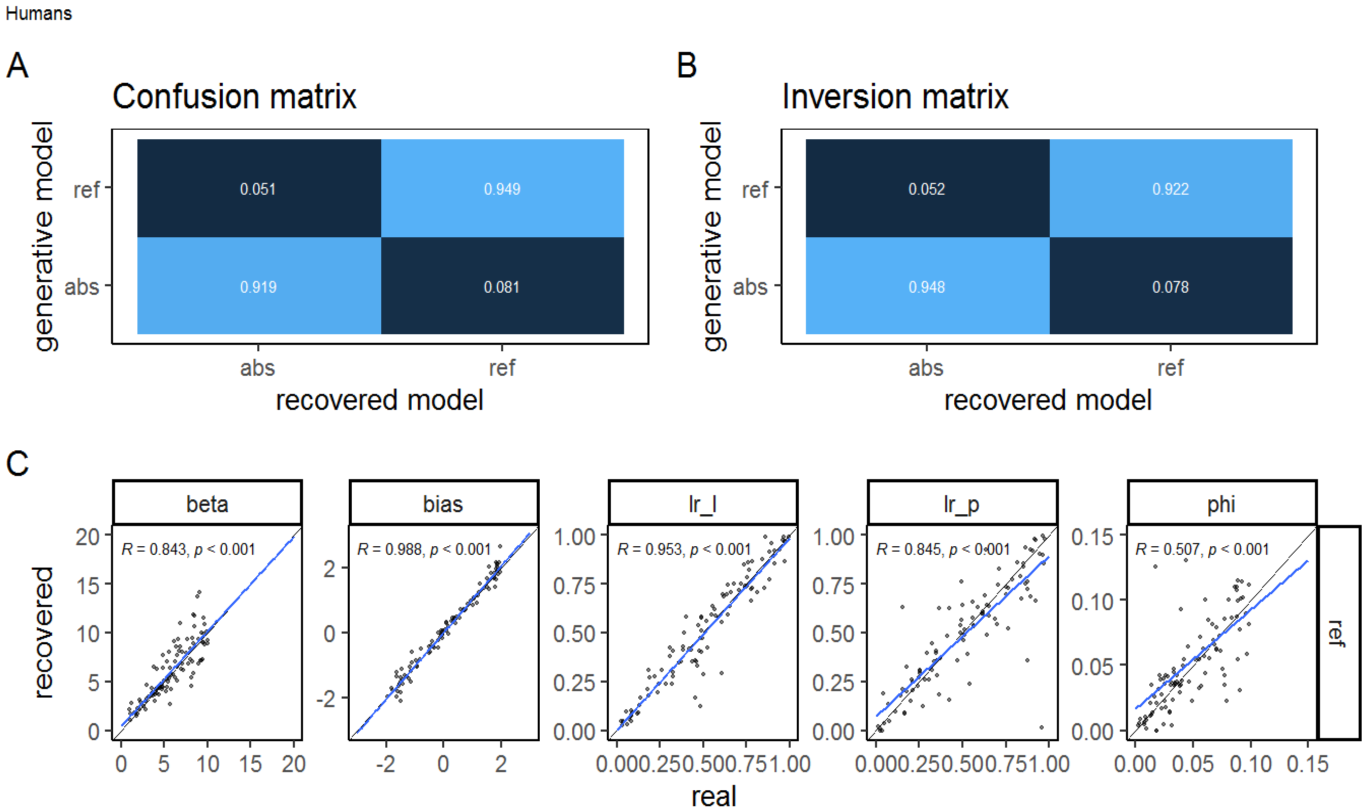


**Supplemental Figure 4: Model and parameter recovery – humans**. "abs" refers to the "Absolute" model, while "ref" refers to the "Reference" model. Parameters lr_l and lr_p refer to the learning rates for the learning and transfer phase respectively, while bias reflects motor bias for options presented on the left and phi is the decay parameter governing the rate of forgetting.

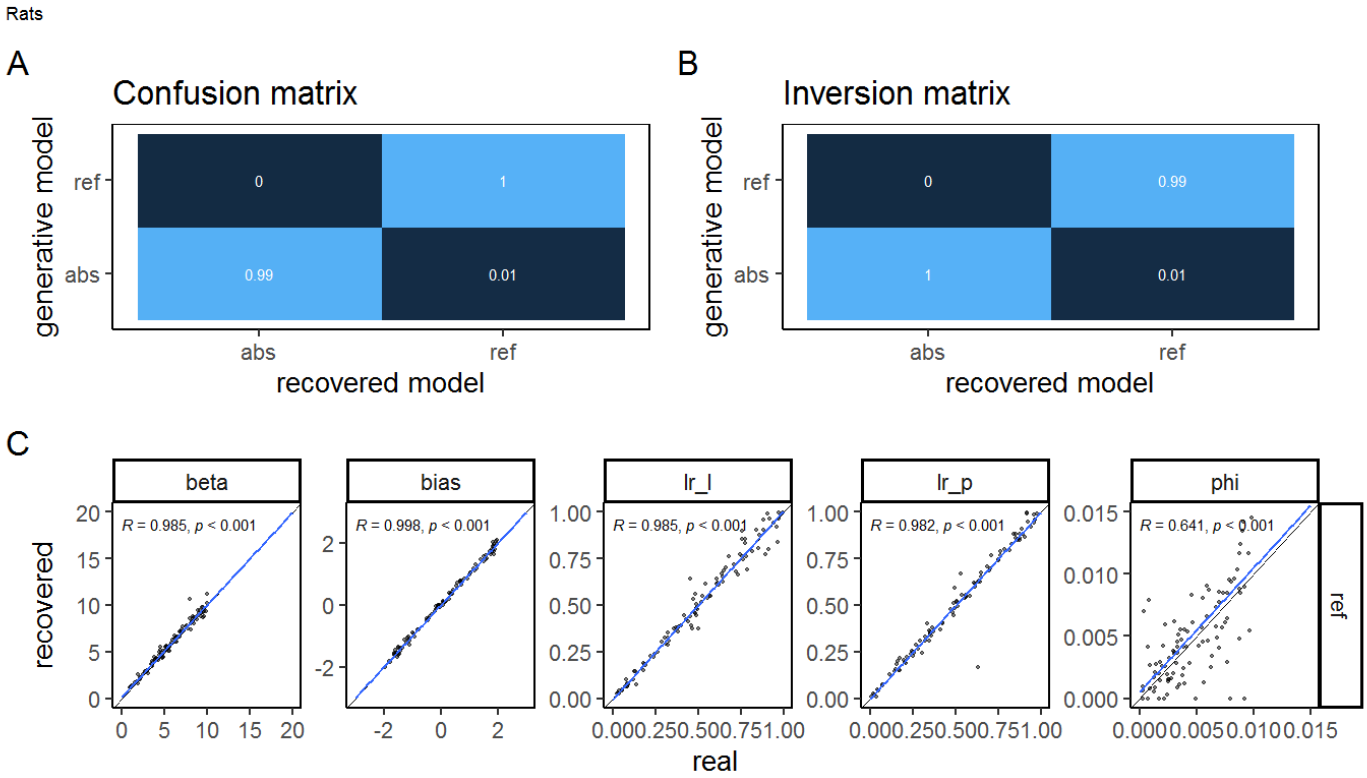


**Supplemental Figure 5: Model and parameter recovery – rats**. "abs" refers to the "Absolute" model, while "ref" refers to the "Reference" model. Parameters lr_l and lr_p refer to the learning rates for the learning and transfer phase respectively, while bias reflects motor bias for options presented on the left and phi is the decay parameter governing the rate of forgetting.

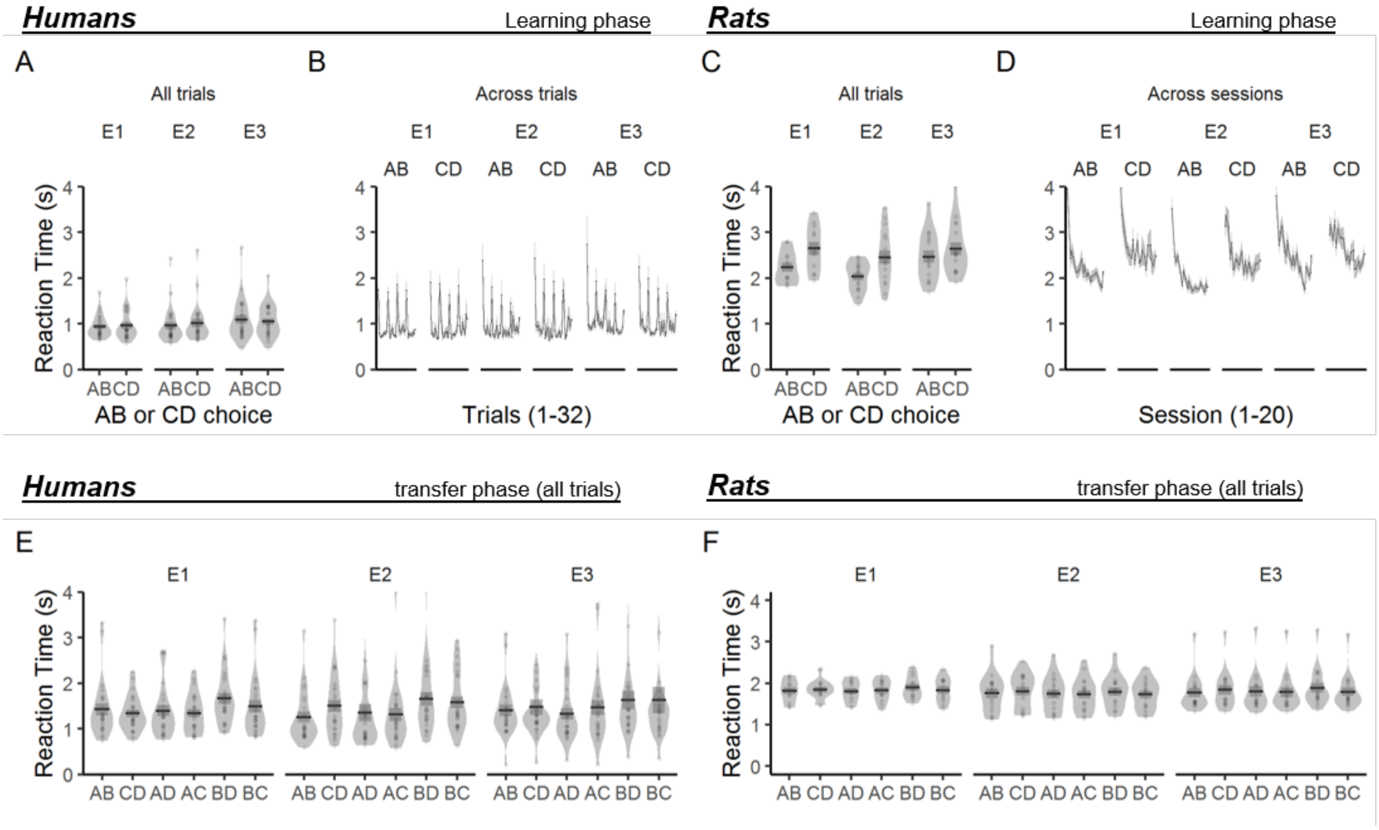


**Supplemental Figure 6:** Reaction times. (**A-B**) Human reaction times in the learning phase. The spikes in B represent start of miniblocks with the same choice pair. (**C-D**) Rat reaction times in the learning phase. (**E**) Human reaction time in the transfer phase. (**F**) Rat reaction times in the transfer phase.

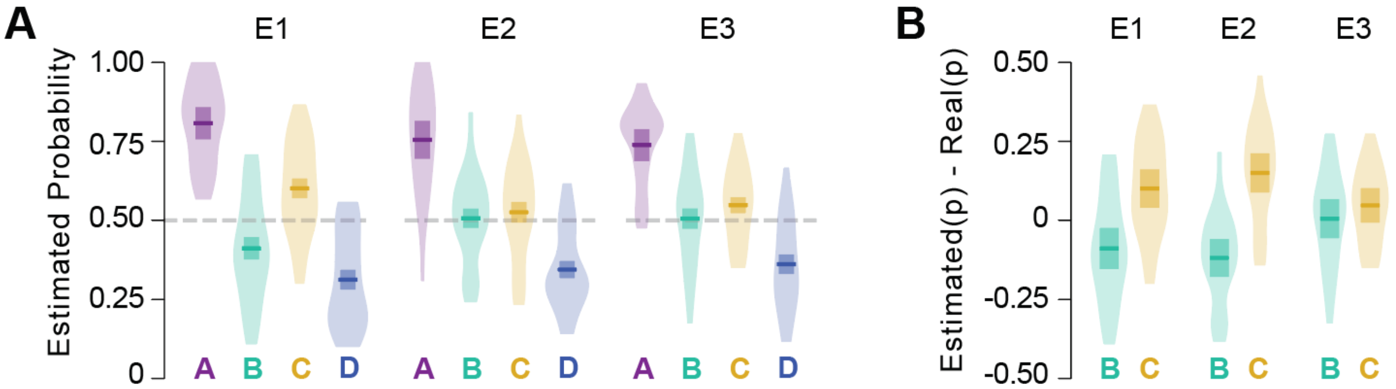


**Supplemental Figure 7: Human participant's explicit rating of reward probability.** (A) Explicit rating of reward probability for each option. (B) Explicit rating of reward probability for critical BC choice options, minus their true probability. Plots clearly show participants' propensity to overestimate the reward probability of C, which was advantageous in the learning phase compared to B which was disadvantageous. Note that these ratings were taken after the end of the transfer phase. The crossbar represents mean and s.e.m.

#### Supplemental Tables

**Supplementary Table 1**. Estimated marginal means of accuracy in the learning phase. All are significantly above chance level (0.5). Note that accuracy the learning phase for rats is calculated in the unrewarded "probe" trials (see **Methods** and **Figure 1**)

| | | **Humans** | | **Rats** | |
|---|---|---|---|---|---|
| | | **Average** | **Slope** | **Average** | **Slope** |
| AB | **E1** | 0.978 [0.959 , 0.988]<br>z=11.6, p<0.001 | 0.068 [0.048, 0.089]<br>z=6.54, p<0.001 | 0.714, [0.66 , 0.761]<br>z=7.22, p<0.001 | 0.036 [0.011, 0.061]<br>z=2.79, p=0.005 |
| | **E2** | 0.948 [0.91 , 0.971]<br>z=9.66, p<0.001 | 0.066 [0.048, 0.085]<br>z=7.10, p<0.001 | 0.784, [0.738 , 0.824]<br>z=10.1, p<0.001 | 0.042 [0.020, 0.064]<br>z=3.80, p<0.001 |
| | **E3** | 0.918 [0.863 , 0.953]<br>z=8.16, p<0.001 | 0.065 [0.047, 0.083]<br>z=7.13, p<0.001 | 0.676, [0.628 , 0.721]<br>z=6.75, p<0.001 | 0.046 [0.024 , 0.068]<br>z=4.17, p<0.001 |
| CD | **E1** | 0.985 [0.971 , 0.992]<br>z=12.3, p<0.001 | 0.103 [0.082, 0.124]<br>z=9.64, p<0.001 | 0.746, [0.703 , 0.785]<br>z=9.74, p<0.001 | 0.079 [0.053 , 0.105]<br>z=5.86, p<0.001 |
| | **E2** | 0.919 [0.864, 0.953]<br>z=8.22, p<0.001 | 0.049 [0.031, 0.067]<br>z=5.37, p<0.001 | 0.675, [0.627 , 0.720]<br>z=6.73, p<0.001 | 0.123 [0.101 , 0.146]<br>z=10.7, p<0.001 |
| | **E3** | 0.962 [0.932 , 0.979]<br>z=10.5, p<0.001 | 0.060 [0.041, 0.079]<br>z=6.25, p<0.001 | 0.688, [0.64 , 0.732]<br>z=7.22, p<0.001 | 0.066 [0.045 , 0.088]<br>z=5.98, p<0.001 |

**Supplementary Table 2**. Estimated marginal means of the accuracy in the transfer phase in the first trial and across all trials, and the estimated increase in accuracy (slope) per trial. Slope estimates were backtransformed from the logarithmic scale. Statistically significant estimates in bold.

| | | Humans | | |
|---|---|---|---|---|
| | | First Trial | All trials | Slope |
| AB | E1 | **0.890 [0.804 , 0.941] z=6.07, p<0.001** | **0.866, [0.815 0.904] z=9.57, p<0.001** | -0.027 [-0.099, 0.044] z=-0.743, p=0.457 |
| | E2 | **0.848 [0.754 , 0.911] z=5.6, p<0.001** | **0.824, [0.765 0.871] z=8.27, p<0.001** | -0.021 [-0.086, 0.043] z=-0.642, p=0.521 |
| | E3 | **0.827 [0.729, 0.895] z=5.31, p<0.001** | **0.793, [0.728 0.846] z=7.34, p<0.001** | -0.027 [-0.088, 0.035] z=-0.848, p=0.397 |
| CD | E1 | **0.887 [0.804, 0.937] z=6.22, p<0.001** | **0.823, [0.764 0.871] z=8.22, p<0.001** | -0.062 [-0.130, 0.005] z=-1.81, p=0.071 |
| | E2 | **0.705 [0.591, 0.799] z=3.39, p<0.001** | **0.695, [0.617 0.763] z=4.67, p<0.001** | -0.006 [-0.061, 0.049] z=-0.208, p=0.835 |
| | E3 | **0.787 [0.683, 0.864] z=4.73, p<0.001** | **0.74, [0.667 0.801] z=5.85, p<0.001** | -0.032 [-0.090, 0.026] z=-1.07, p=0.284 |
| AD | E1 | **0.859 [0.763, 0.92] z=5.56, p<0.001** | **0.871, [0.821, 0.908] z=9.76, p<0.001** | 0.012 [-0.057, 0.082] z=0.352, p=0.725 |
| | E2 | **0.844 [0.748, 0.908] z=5.5, p<0.001** | **0.832, [0.774, 0.878] z=8.52, p<0.001** | -0.010 [-0.075, 0.054] z=-0.312, p=0.755 |
| | E3 | **0.763 [0.65, 0.849] z=4.14, p<0.001** | **0.846, [0.791, 0.889] z=8.94, p<0.001** | **0.064 [0.002, 0.127] z=2.02, p=0.043** |
| AC | E1 | 0.607 [0.488, 0.715] z=1.77, p=0.077 | **0.693, [0.616, 0.761] z=4.64, p<0.001** | 0.046, [-0.007, 0.099] z=1.69, p=0.092 |
| | E2 | **0.635 [0.515, 0.74] z=2.2, p=0.028** | **0.723, [0.648, 0.787] z=5.38, p<0.001** | 0.049, [-0.006, 0.103] z=1.76, p=0.078 |
| | E3 | 0.532 [0.413, 0.647] z=0.525, p=0.600 | **0.705, [0.628, 0.772] z=4.92, p<0.001** | 0.090 [0.036, 0.143] z=3.29, p=0.001 |
| BD | E1 | 0.477 [0.363, 0.594] z=-0.381, p=0.703 | **0.61, [0.527, 0.687] z=2.57, p=0.010** | **0.065 [0.013, 0.116] z=2.46, p=0.014** |
| | E2 | 0.473 [0.358, 0.591] z=-0.449, p=0.65 | **0.605, [0.521, 0.683] z=2.44, p=0.015** | **0.065 [0.012, 0.117] z=2.42, p=0.015** |
| | E3 | **0.621 [0.502, 0.726] z=1.99, p=0.046** | **0.661, [0.58, 0.733] z=3.82, p<0.001** | 0.021 [-0.032, 0.074] z=0.785, p=0.433 |
| BC | E1 | **0.173 [0.106, 0.269] z=-5.43, p<0.001** | **0.282, [0.217, 0.358] z=-5.23, p<0.001** | **0.076 [0.017, 0.136] z=2.51, p=0.012** |
| | E2 | **0.272 [0.184, 0.382] z=-3.82, p<0.001** | 0.463, [0.38, 0.548] z=-0.846, p=0.398 | **0.101 [0.046, 0.155] z=3.63, p<0.001** |
| | E3 | **0.299 [0.206, 0.411] z=-3.38, p<0.001** | 0.439, [0.358, 0.524] z=-1.41, p=0.159 | **0.073, [0.020,0.127] z=2.71, p=0.007** |

**Supplementary Table 3**. Estimated marginal means of the accuracy in the transfer phase in the first session and across all session, and the estimated increase in accuracy (slope) per session. Statistically significant estimates in bold.

| | | Rats | | |
|---|---|---|---|---|
| | | First Session | All Sessions | Slope |
| AB | E1 | **0.754 [0.692 , 0.806] z=7.12, p<0.001** | **0.779, [0.725 0.824] z=8.6, p<0.001** | 0.031 [-0.009 , 0.071] z=1.52, p=0.130 |
| | E2 | **0.662 [0.602 , 0.717] z=5.1, p<0.001** | **0.719, [0.667 0.766] z=7.51, p<0.001** | **0.060, [0.027, 0.092] z=3.6, p<0.001** |
| | E3 | **0.694 [0.637, 0.746, z=6.21, p<0.001** | **0.704, [0.65 0.752] z=6.91, p<0.001** | 0.010 [-0.02, 0.042] z=0.606, p=0.545 |
| CD | E1 | **0.815 [0.763, 0.858] z=9.23, p<0.001** | **0.778, [0.725 0.824] z=8.58, p<0.001** | **-0.051 [-0.091, -0.011] z=-2.51, p=0.012** |
| | E2 | **0.809 [0.764, 0.848] z=10.4, p<0.001** | **0.774, [0.727 0.814] z=9.72, p<0.001** | **-0.048 [-0.082, -0.014] z=-2.73, p=0.006** |
| | E3 | **0.720 [0.665, 0.769] z=7.14, p<0.001** | **0.687, [0.632 0.737] z=6.3, p<0.001** | **-0.035 [-0.067, -0.004] z=-2.2, p=0.028** |
| AD | E1 | **0.836 [0.786, 0.876] z=9.79, p<0.001** | **0.841, [0.798, 0.876] z=11.2, p<0.001** | 0.008, [-0.036, 0.052] z=0.348, p=0.728 |
| | E2 | **0.843 [0.802, 0.878] z=11.5, p<0.001** | **0.866, [0.834, 0.893] z=14.3, p<0.001** | **0.041, [0.002, 0.081] z=1.97, p=0.049** |
| | E3 | **0.771 [0.72, 0.815] z=8.87, p<0.001** | **0.783, [0.738, 0.822] z=10.1, p<0.001** | 0.016, [-0.019, 0.050] z=0.887, p=0.375 |
| AC | E1 | 0.481 [0.408, 0.553] z=-0.523, p=0.601 | **0.584, [0.514, 0.65] z=2.36, p=0.018** | **0.093, [0.057, 0.128] z=5.15, p<0.001** |
| | E2 | 0.498 [0.435, 0.561] z=-0.0645, p=0.949 | **0.616, [0.557, 0.672] z=3.8, p<0.001** | **0.107, [0.076, 0.138] z=6.77, p<0.001** |
| | E3 | 0.544 [0.482, 0.606] z=1.39, p=0.166 | **0.575, [0.515, 0.633] z=2.43, p=0.015** | 0.027 [-0.003, 0.057] z=1.79, p=0.073 |
| BD | E1 | **0.675 [0.607, 0.736] z=4.82, p<0.001** | **0.628, [0.56, 0.691] z=3.65, p<0.001** | **-0.046 [-0.081, -0.010] z=-2.52, p=0.012** |
| | E2 | **0.693 [0.635, 0.745] z=6.16, p<0.001** | **0.681, [0.626, 0.732] z=6.08, p<0.001** | -0.012 [-0.043, 0.020] z=-0.743, p=0.458 |
| | E3 | **0.571 [0.508, 0.631] z=2.21, p=0.027** | **0.577, [0.517, 0.635] z=2.5, p=0.012** | 0.006 [-0.024, 0.036] z=0.376, p=0.707 |
| BC | E1 | **0.304 [0.244, 0.371] z=-5.42, p<0.001** | **0.302, [0.245, 0.364] z=-5.81, p<0.001** | -0.002 [-0.039, 0.037] z=-0.122, p=0.903 |
| | E2 | **0.332 [0.278, 0.392] z=-5.33, p<0.001** | **0.387, [0.331, 0.446] z=-3.7, p<0.001** | **0.053, [0.022, 0.083] z=3.36, p<0.001** |
| | E3 | **0.358 [0.301, 0.418] z=-4.51, p<0.001** | **0.374 [0.319, 0.433] z=-4.14, p<0.001** | 0.016 [-0.015, 0.046] z=1.01, p=0.312 |

**Supplementary Table 4.** Demographic data. Mean and standard deviation of sex, age, bonus payment and completion time across all experiments.

| | n | Sex | | Age | Bonus Payment | Time |
|---|---|---|---|---|---|---|
| | | M | F | | | |
| **E1** | 24 | 12 | 12 | 26.5 ± 6.80 | 2.10 ± 0.128 | 23.9 ± 8.85 |
| **E2** | 24 | 12 | 12 | 30.1 ± 10.7 | 1.91 ± 0.199 | 24.1 ± 7.75 |
| **E3** | 24 | 12 | 12 | 33.0 ± 12.9 | 1.94 ± 0.154 | 23.2 ± 7.17 |

**Supplementary Table 5.** Stimulus features

| Stimulus | Orientation | Frequency | Background Colour (RGB) | Subjects |
|---|---|---|---|---|
| 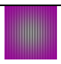 | 0 | 0.2 | 153, 0, 153 | Humans & Rats |
| 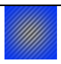 | 45 | 0.15 | 0, 51, 255 | Humans & Rats |
| 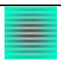 | 90 | 0.1 | 0, 255, 204 | Humans & Rats |
| 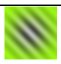 | 135 | 0.05 | 153, 255, 0 | Humans & Rats |
| 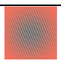 | 22.5 | 0.25 | 255, 113, 91 | Humans |
| 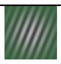 | 67.5 | 0.075 | 56, 102, 65 | Humans |
| 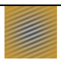 | 112.5 | 0.125 | 200, 153, 51 | Humans |
| 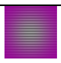 | 157.5 | 0.175 | 200, 121, 255 | Humans |

**Supplementary Table 6**. Estimated marginal means of reaction times in the learning phase.

| | | Humans | Rats |
|---|---|---|---|
| **AB** | **E1** | M=787, CI=[712;871], | M=2030, CI=[1830;2250], |
| | **E2** | M=769, CI=[696;851], | M=1810, CI=[1650;1980], |
| | **E3** | M=875, CI=[791;968], | M=2160, CI=[1970;2360], |
| **CD** | **E1** | M=762, CI=[689;843], | M=2270, CI=[2050;2520], |
| | **E2** | M=767, CI=[693;848], | M=2110, CI=[1930;2310], |
| | **E3** | M=839, CI=[758;928], | M=2320, CI=[2120;2530], |

**Supplementary Table 7**. Estimated marginal means of reaction times in the transfer phase.

| | | Humans | Rats |
|---|---|---|---|
| **AB** | **E1** | M=1040, CI=[906;1180] | AB M=1660, CI=[1490;1840] |
| | **E2** | M=1000, CI=[876;1150] | AB M=1560, CI=[1430;1710] |
| | **E3** | M=1080, CI=[947;1240] | AB M=1630, CI=[1490;1780] |
| **CD** | **E1** | M=1070, CI=[939;1230] | CD M=1680, CI=[1510;1860] |
| | **E2** | M=1130, CI=[991;1300] | CD M=1590, CI=[1450;1740] |
| | **E3** | M=1160, CI=[1010;1330] | M=1680, CI=[1540;1840] |
| **AD** | **E1** | M=1020, CI=[894;1170] | M=1670, CI=[1500;1850] |
| | **E2** | M=1010, CI=[887;1160] | M=1570, CI=[1430;1710] |
| | **E3** | M=1040, CI=[909;1190] | M=1630, CI=[1490;1790] |
| **AC** | **E1** | M=1050, CI=[921;1200] | M=1670, CI=[1510;1860] |
| | **E2** | M=1030, CI=[899;1180] | M=1560, CI=[1420;1700] |
| | **E3** | M=1120, CI=[980;1280] | M=1620, CI=[1480;1780] |
| **BD** | **E1** | M=1310, CI=[1150;1500] | M=1730, CI=[1560;1920] |
| | **E2** | M=1240, CI=[1080;1420] | M=1630, CI=[1490;1780] |
| | **E3** | M=1250, CI=[1090;1430] | M=1720, CI=[1570;1880] |
| **BC** | **E1** | M=1130, CI=[990;1290] | M=1700, CI=[1530;1890] |
| | **E2** | M=1240, CI=[1080;1410] | M=1560, CI=[1430;1710] |
| | **E3** | M=1150, CI=[1010;1320] | M=1660, CI=[1520;1820] |
